# β3 accelerates microtubule plus end maturation through a divergent lateral interface

**DOI:** 10.1101/2024.07.17.603993

**Authors:** Lisa M. Wood, Jeffrey K. Moore

**Affiliations:** Department of Cell and Developmental Biology, University of Colorado Anschutz Medical Campus, Aurora, CO, USA

## Abstract

β-tubulin isotypes exhibit similar sequences but different activities, suggesting that limited sequence divergence is functionally important. We investigated this hypothesis for TUBB3/β3, a β-tubulin linked to aggressive cancers and chemoresistance in humans. We created mutant yeast strains with β-tubulin alleles that mimic variant residues in β3 and find that residues at the lateral interface are sufficient to alter microtubule dynamics and response to microtubule targeting agents. In HeLa cells, β3 overexpression decreases the lifetime of microtubule growth, and this requires residues at the lateral interface. These microtubules exhibit a shorter region of EB binding at the plus end, suggesting faster lattice maturation, and resist stabilization by paclitaxel. Resistance requires the H1-S2 and H2-S3 regions at the lateral interface of β3. Our results identify the mechanistic origins of the unique activity of β3 tubulin and suggest that tubulin isotype expression may tune the rate of lattice maturation at growing microtubule plus ends in cells.

## Introduction

Assembling a microtubule requires heterodimeric αβ-tubulin subunits that bind neighboring subunits along longitudinal and lateral sides and modulate those interactions through a GTPase cycle. Heterodimers bound to GTP preferentially incorporate at the growing plus end of the microtubule, stimulating the GTP hydrolysis cycle that subsequently powers a wave of conformational changes across the heterodimer that rearrange binding interfaces and alter the stability of the microtubule lattice. This creates zones in the microtubule that are distinguished by the nucleotide status and stability of the resident heterodimers (Alushin et al., 2014, Geyer et al., 2015, Hyman et al., 1995). Starting from the terminal plus end, the ‘GTP cap’ is a structurally heterogeneous region where newly incorporated GTP-bound heterodimers rearrange from the bent conformation they exhibit in solution to a straight conformation that supports lateral interactions with neighboring protofilaments and completes the catalytic pocket for GTP hydrolysis (McIntosh et al., 2018, Nogales et al., 1998, Nogales et al., 1999, Mickolajczyk et al., 2019, Carlier and Pantaloni, 1981). The ‘transition zone’ is the region of GTP hydrolysis where heterodimers are bound to GDP•Pi and exhibit a shift in protofilament skew that suggests a rearrangement of lateral interactions (Estevez-Gallego et al., 2020, Manka and Moores, 2018). This region is recognized by end-binding (EB) proteins that act as regulators of microtubule dynamics and target many proteins to growing microtubule ends (Zanic et al., 2009, Maurer et al., 2014, Honnappa et al., 2009). Finally, the ‘GDP-lattice’ region contains subunits that have released γ-phosphate. Heterodimers in this region create an unstable lattice with weaker lateral interactions and stronger longitudinal interactions, suggesting a mechanism through which the energy from GTP hydrolysis could be stored in the lattice to eventually drive the outward peeling of protofilaments when the lattice disassembles (Mandelkow et al.,1991, Zhang et al., 2015, Manka and Moores, 2018).

Microtubule dynamics, the ability of microtubules to switch between polymerization and depolymerization, underlies microtubule function in a wide array of cellular process including mitosis, migration, and organelle transport. The requirement for microtubule dynamics in these processes makes an attractive target for therapeutics. Indeed, Microtubule-Targeting Agents (MTAs) are effective therapeutics for cancers and other diseases, and generally act by preventing αβ-heterodimers from transiting into different conformational states and zones of the microtubule. For example, colchicine binds to soluble heterodimers, vinca alkaloids bind heterodimers at the ends of microtubules, and paclitaxel binds heterodimers within the lumen of a formed microtubule (Bai et al., 1990, Uppuluri et al., 1993, Nogales et al., 1998). These serve to target curved GTP-tubulin in solution, the GTP-cap, and GDP-lattice, respectively. MTAs have had clinical success, however tumors frequently become refractory to treatment, or patients experience severe side effects due to the widespread reliance of cells on dynamic microtubules. Therefore, understanding the mechanisms that control heterodimer interactions and transit between zones could improve the design of MTAs and limit the potential for drug resistance.

MTA resistance may be partly attributed to the heterogeneity of αβ-tubulin proteins. Most eukaryotes express families of genes for αβ-tubulin, known as tubulin isotypes. Humans have 9 α and 8 β tubulin isotypes which are distinct genes with expression patterns that differ across cell type and developmental stage (reviewed in Gasic, 2022). Furthermore, isotypes code for tubulins with distinct amino acid sequences and, in some cases, biochemical activities. The class 3 β-tubulin isotype, also known as β3 or TUBB3 in humans, is primarily expressed in young neurons during the early stages of neurite outgrowth and decreases in expression as the neurons mature (Hausrat et al., 2021, Park et al., 2021). Missense mutations in *TUBB3* are associated with a variety of neurodevelopmental disorders (Tischfield et al., 2010, reviewed in Breuss et al., 2017). It is also well-established that *TUBB3* expression increases across several cancers including breast and lung cancers, and correlates with poor prognosis and drug-resistant cancer (Seve et al., 2007, Bernard-Marty et al., 2002, Ferrandina et al., 2006). Due to its association with disease, β3 has been studied in reconstitution experiments to examine its biochemical activity. These studies consistently show that microtubules assembled from heterodimers containing β3 exhibit a higher frequency of catastrophe, the switch from polymerization to depolymerization, and some studies show an increased rate of depolymerization (Panda et al., 1994, Pamula et al., 2016, Ti et al., 2018, Cross and Chew, 2023, Vemu et al., 2016). These observations have led to a model where β3 increases microtubule dynamics and promotes a more rapid turnover of microtubules. In addition, β3 has been implicated in resistance to the MTA paclitaxel, which stabilizes the microtubule lattice. (Parker et al., 2017, Stengel et al., 2010, Chew and Cross, 2023)

What remains to be understood is the mechanism through which β3 alters interactions between heterodimers and the transit between zones in a microtubule. The domains of β-tubulin that are responsible for increasing catastrophe frequency are unknown and the mechanism through which β3 promotes resistance to paclitaxel remains to be investigated. We sought to answer these questions through a combination of modeling β3 in *S. cerevisiae* and quantitative imaging of HeLa cells expressing β-tubulin mutants. We identify sequence variants in β3 that are sufficient to alter microtubule dynamics in a homogenous tubulin pool and furthermore show that sequence variants in the H1-S2 and H2-S3 regions are necessary for the unique effects of β3 on microtubule dynamics and paclitaxel response in human cells.

## Results

### Modeling β3 residues in yeast β-tubulin

TUBB3/β3 is a divergent isotype (Figure 1A). Whereas five of the human β-tubulin isotypes are closely related and exhibit greater than 95% sequence identity compared to each other, three other isotypes – TUBB1/β1, TUBB3/β3 and TUBB6/β6 – exhibit less than 95% identity to any other isotype (Figure 1B). Amongst these three, β3 and β6 are most closely related to each other, and β1 is the most divergent. Sequence alignment comparing β3 to the other β-tubulin isotypes identified 33 residues that differ from at least three of the other human β-tubulin isotypes at the same position (Figure S1; table 1). In addition, the CTT of β3 is five residues longer than most other human β-tubulin isotypes. We will refer to these collectively as β3 variant residues. We reasoned that β3 variant residues are likely to contribute to differences in tubulin activity.

**Figure 1:**
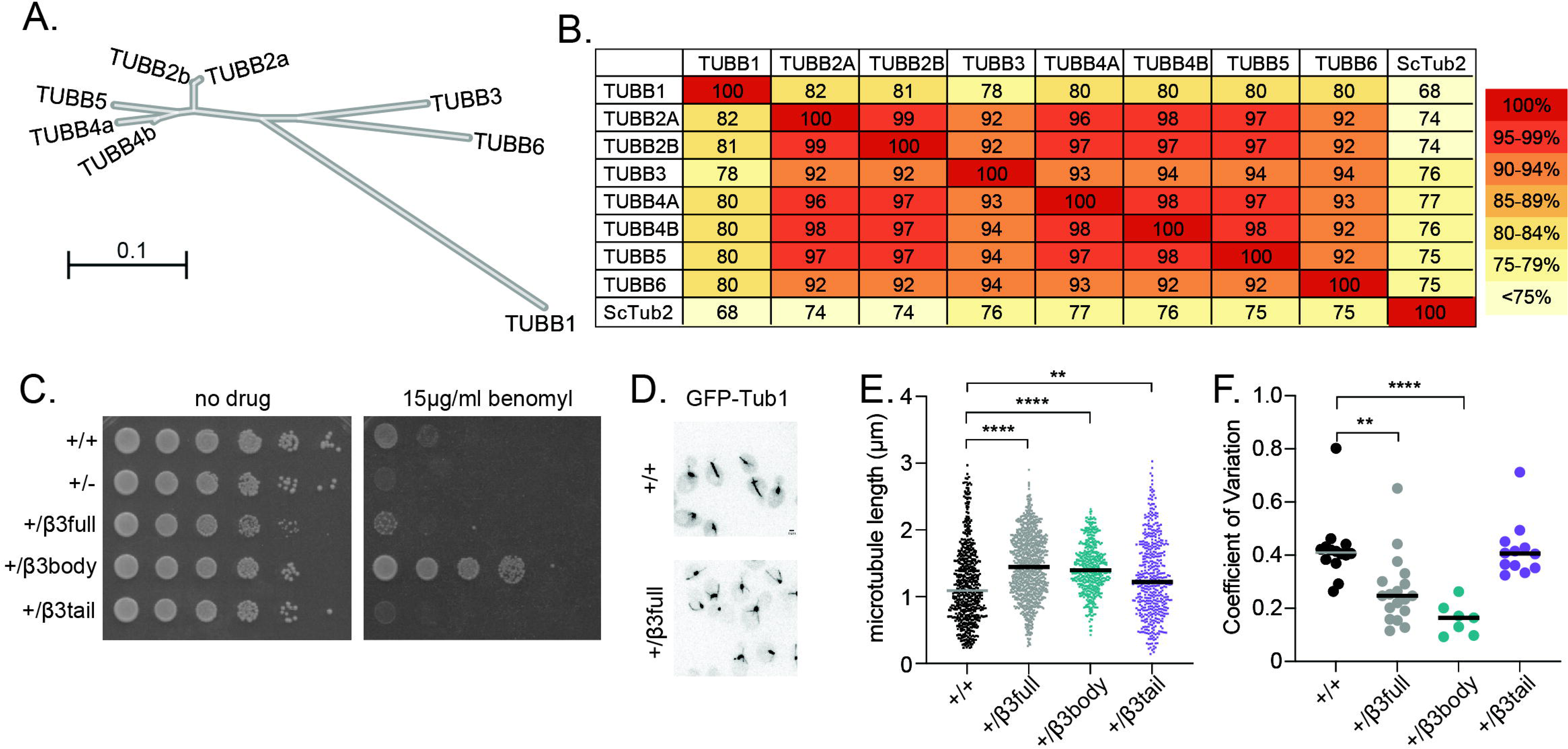
β3 variant residues impart microtubule stabilization in *S. cerevisiae*. **(A)** Radial phylogenetic tree of 8 *H. sapiens* β-tubulin isotypes generated using Clustal Omega (Madiera et al., 2024). Scalebar is the average amino acid variance per residue. **(B)** Amino acid alignment of *H. sapiens* β-tubulins with percent identity reported. The last row/column shows *S.c* β-tubulin aligned with the various *H.s* β-tubulins. **(C)** 10-fold serial dilution of diploid β3-mimic strains indicated at left. Cells were spotted to rich medium or rich medium supplemented with 15 µg/ml benomyl and grown for 2 days at 30°C. **(D)** Representative field of wild-type or β3full diploid cells expressing GFP-Tub1. **(E)** Astral microtubule lengths (µm) from timelapse imaging collected at 5-second intervals for 5 minutes, for the indicated genotypes. At least 14 cells were analyzed for each genotype. P-values are as follows; ** = 0.0094, ***** <0.0001. **(F)** Coefficient of variation of each microtubule in our timelapse imaging was calculated by dividing the standard deviation by the mean. Each dot represents one microtubule. P-values are as follows; **=0.0021, ****<0.0001.

**Table 1.** Tubb3 variant residues.

| Region | Functional domain | Residue | *Hs* TUBB3 | Consensus<br>*Hs* β-isotype | *Sc* Tub2 |
| --- | --- | --- | --- | --- | --- |
| H1-H1' | Lateral interface | 33 | S | T | N |
| H1-H1' | Lateral interface | 37 | V | H | H |
| H1' | Lateral interface | 48 | S | N | N |
| H1'-S2 | Lateral interface | 56 | S | G | S |
| H1'-S2 | Lateral interface | 57 | H | N | G |
| H2-H2' | Lateral interface | 80 | A | P | A |
| H2'-H2'' | Lateral interface | 83 | H | Q | N |
| H2'-H2'' | Lateral interface | 84 | L | I | L |
| S3 | Lateral interface | 91 | I | V | I |
| H3 | Core | 126 | N | S | G |
| H4 | Core | 155 | V | I | I |
| T5 loop | Longitudinal interface<br>and GTP pocket | 170 | V | M | L |
| H5 | Core | 189 | I | V | V |
| H6-H7 | Core | 218 | A | T | N |
| H7 | Core | 239 | S | C | S |
| M loop | Lateral interface and<br>ptx binding pocket | 275 | A | S | A |
| S8 | Core | 315 | T | A | A |
| H10 | Core | 333 | I | V | V |
| S9 | Core | 351 | V | T | T |
| S10 | Core | 365 | S | A | A |

To investigate the impact of β3 variant residues we created a mimetic allele using the budding yeast β-tubulin *TUB2*. Native Tub2 protein already contains 6 of the β3 variant residues (Table 1), therefore we created 14 substitutions in the globular body of Tub2 to mimic β3, and replaced the Tub2 CTT with the 20 amino acid CTT from β3. We refer to this allele as ‘β3full’. Heterozygous diploid yeast expressing one copy of β3full and one copy of wild-type *TUB2* exhibit slightly slower growth on rich media, compared to wild-type controls and hemizygous diploids that express only one copy of wild-type *TUB2* (Figure 1C; ‘no drug’ panel). When challenged with the microtubule destabilizing agent, benomyl, cells expressing β3full display an intermediate phenotype that is more sensitive than wild-type but slightly more resistant than the hemizygote (Figure 1C). This indicates that β3full is not equivalent to wild-type Tub2, but does provide some weakened level of β-tubulin function in yeast.

To determine how β3full effects microtubule dynamics, we integrated two copies of ectopically expressed GFP-Tub1 into the β3full heterozygous diploid and the wild-type diploid. By observing microtubules over time in living cells, we noted several differences for β3full. Microtubules in β3full cells are long and curl around the cell cortex, and we occasionally observe microtubules that break off to create fragments that persist for up to minutes in the cell (Figure 1D; Video S1). By measuring the lengths of astral microtubules in populations of cells over time, we find that microtubules in β3full cells are significantly longer than in wild-type cells and the lengths of individual microtubules exhibit a smaller coefficient of variation (Figure 1E and F). This suggests that microtubules containing β3full tubulin are more stable than wild-type microtubules. This was surprising, since purified mammalian heterodimers containing β3 have been shown to increase catastrophe frequency compared to other isotypes (Vemu et al., 2016, Pamula et al., 2016, Chew and Cross, 2023). Our results suggest that in the context of yeast tubulin, the β3 variant residues may stabilize microtubules.

We next asked whether the effects of the β3full allele could be attributed to variant residues in the globular domain of β-tubulin or in the CTT. We created two alleles to separate the globular domain from the CTT; ‘β3body’ contains 13 β3 variant residues within the globular domain of Tub2, and ‘β3tail’ contains the native Tub2 globular domain and the CTT of β3 (Table 1). Heterozygous diploids expressing one copy of β3body or β3tail and one copy of wild-type *TUB2* exhibit wild-type levels of growth on rich media, but show divergent phenotypes when challenged with benomyl (Figure 1C). β3body cells show increased benomyl resistance compared to wild-type controls, while β3tail cells show increased benomyl sensitivity. These alleles also show divergent phenotypes in our microscopy assay imaging GFP-labeled microtubules. Microtubules in β3body cells are longer than wild-type controls and exhibit a decreased coefficient of variation, but are not significantly different from β3full (Figure 1E and F). We did not observe microtubules breaking and creating fragments in β3body cells (Video S2). In contrast, microtubules in β3tail cells are significantly shorter than in β3full cells, and exhibit an increased coefficient of variation (Figure 1E and F). These β3tail microtubules are more similar to wild-type controls, but slightly longer (Figure 1E). These results suggest that β3 variant residues within the globular domain are responsible for the stabilizing effects seen in the β3full allele, and that the CTT of β3 may mitigate the effects of the globular domain variants.

### Identifying β3 variant residues that alter responses to MTAs

We next sought to determine the contributions of the individual β3 variant residues to β-tubulin activity. Using previously published structural analysis, we identified TUBB3 variant residues positioned at lateral and longitudinal interfaces, nucleotide binding pockets, and the paclitaxel binding pocket (Figure 2A and B; Figure S2). All other residues are buried within the globular domain of β-tubulin, and categorized as ‘core’. Domain enrichment analysis shows that β3 variant residues are statistically enriched at the lateral interface, while no enrichment was detected at the longitudinal interface, GTP binding pocket, or paclitaxel binding pocket (Figure 2A; p=0.024 for lateral interface). We also find that variant residues are enriched in the CTT (Figure 2A; p=0.003; Figure S1). These data indicate that β3 variant residues are enriched in domains of β-tubulin where they may affect interactions with neighboring protofilaments or interactions at the microtubule surface.

**Figure 2:**
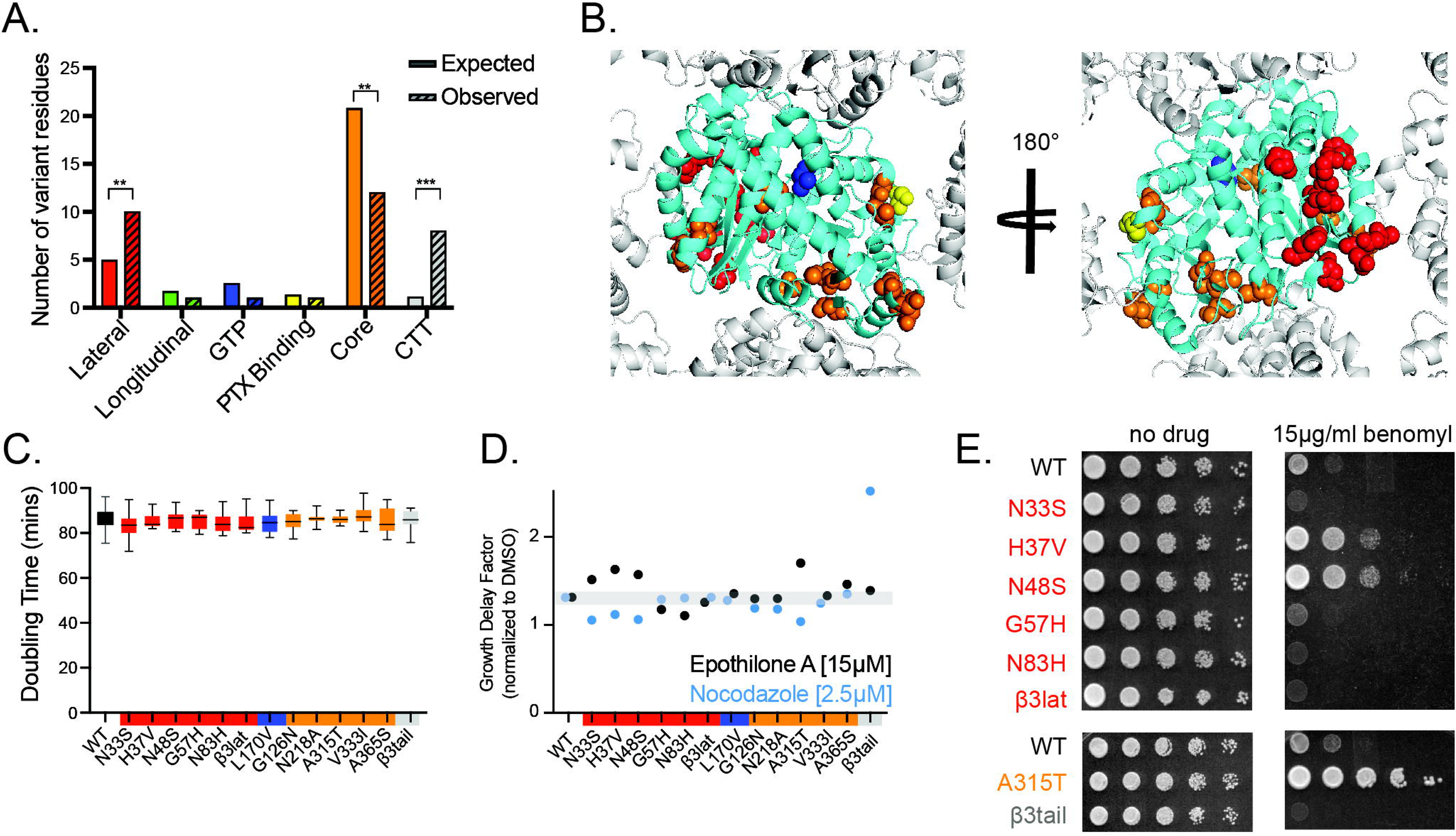
Individual β3 variant residues modulate response to microtubule targeting agents. **(A)** TUBB3 variant residues categorized by functional domain: lateral interface, longitudinal interface, GTP-binding pocket, paclitaxel (PTX) binding pocket, core, or the CTT. Expected number of variant residues was determined by multiplying the proportion of amino acids in each domain by 33 which is the number of β3 variant residues including the CTT. This represents the number of mutations that would be expected from a random distribution throughout the tubulin protein. Observed and expected variant residues were compared using a Chi-squared test to determine if the residues are randomly distributed across domains or enriched. **(B)** Positions of β3 variant residues on β-tubulin contained within a microtubule lattice. The 180° turn displays a view from inside the lumen looking out. Residues are colored according to the domains defined in panel A. **(C)** Liquid growth assay to determine the doubling time of mutants with β3 variant residues. Bars represent mean with 95% CI from triplicate experiments with biological replicates included in each experiment. No significant differences in doubling time were determined. **(D)** Growth delay factor (GDF) of each strain in either 15µM EpoA or 2.5µM Nocodazole. GDF was calculated by dividing the average doubling time in the presence of MTA by the doubling time in DMSO from same experiment. The grey shading represents 95% CI for wild-type GDF in 15µM EpoA and 2.5µM Nocodazole. Values that fell outside the 95% CI were statistically significant and p-values are reported in Figure S3 for the raw doubling times. **(E)** Serial dilution of yeast cells expressing β3 variant residues indicated to the left of the plate. Cells were spotted to rich media or rich media supplemented with 15 µg/ml benomyl and grown for 2 days at 30°C.

We created all the β3 variant residues as single mutations in yeast Tub2. Eleven of the haploid single mutants and the β3tail haploid mutant showed no impact on cellular fitness when grown in rich media (Figure 2C); but we were unable to recover haploid cells with individual mutations at residues 155, 189, and 351. These data indicate that β3 variants at residues 155, 189, and 351 strongly impact β-tubulin function in *S. cerevisiae*, but the majority of β3 variant residues may have subtler impacts on β-tubulin function.

We first examined the impact of β3 variant residues at the lateral interface. The lateral interface associates in a homotypic (β-β) manner during polymerization and is thought to represent the weakest point in the microtubule lattice. We hypothesized that β3 variant residues may alter the homotypic contacts at the lateral interface to modulate the stability of the lattice. To test this, we determined sensitivity or resistance to MTAs that stabilize or destabilize microtubules, Epothilone A (EpoA) and nocodazole, predicting that mutants that destabilize the lattice will exhibit resistance to stabilizing MTAs and hypersensitivity to destabilizing MTAs, and vice versa. We used concentrations of each MTA that significantly limit the growth of wild-type yeast cells, and used the calculated doubling times (Figure S3) to calculate the growth delay factor for each mutant, defined as the fold change compared to a DMSO control for each genotype. Three variant residues at the lateral interface, N33S, H37V, and N48S, show sensitivity to EpoA and resistance to nocodazole, while two other residues at the lateral interface, G57H and N83H, show resistance to EpoA and wild-type levels of sensitivity to nocodazole (Figure 2D). To rule out the possibility that these sensitivities are specific to EpoA or nocodazole, we used a third, destabilizing MTA, benomyl. Two of the nocodazole resistant mutants, H37V and N48S, are also resistant to benomyl, while N33S shows resistance to nocodazole and sensitivity to benomyl (Figure 2E). Additionally, we find that G57H and N83H exhibit slightly increased sensitivity to benomyl, compared to wild-type controls (Figure 2E). These data show that individual β3 variant residues at the lateral interface exhibit distinct sensitivities to MTAs, and that these effects cannot be attributed to altered binding affinity for a particular MTA.

To assess how the five β3 variant residues at the lateral interface work in combination, we created a *TUB2* allele that combined them; we call this allele ‘β3lat’. β3lat cells exhibit a level of sensitivity to EpoA and nocodazole that is similar to wild-type cells, but are hypersensitive to benomyl (Figure 2D and E). This indicates that the combination of TUBB3 variant residues at the lateral interface creates a mildly destabilizing effect that is different from any individual mutant.

We also tested variant residues positioned at other domains. The single variant at the GTP-binding pocket, L170V, shows no changes in sensitivity to EpoA, nocodazole or benomyl (Figure 2D; Figure S3). We conclude that this variant is not sufficient to alter β-tubulin function. Of the five TUBB3 variant residues within the core of the protein, only one showed a significant response in our drug sensitivity assays. A315T is sensitive to EpoA, resistant to nocodazole, and resistant to benomyl (Figure 2D and E). These data indicate that A315T may be stabilize microtubules in yeast. The β3tail allele shows a level of EpoA sensitivity that is similar to wild-type controls, but is hypersensitive to nocodazole and benomyl (Figure 2D and E). These results suggest that the β3 CTT impacts microtubule dynamics differently from variant residues in the β-tubulin body.

In summary, our screen of the β3 variant residues identified several regions that may contribute to the activity of β3: the lateral interface, A315T in the core, and β3tail. Each of these showed differential responses to MTAs in a manner that suggests effects on microtubule dynamics, rather than effects on the binding of specific MTAs.

### Variant residues at the lateral interface and core of β3 are sufficient to alter microtubule dynamics

Differential response to MTAs could indicate differential effects on microtubule dynamics. *In-vitro,* β3 increases catastrophe frequency and, in some studies, depolymerization rate (Pamula et al., 2016, Cross and Chew, 2023, Vemu et al., 2016). We asked whether any of the candidate TUBB3 variant residues is sufficient to create similar changes in microtubule dynamics in yeast. We integrated one copy of GFP-Tub1 into the wild-type, β3lat, A315T, and β3tail haploid strains and measured astral microtubule lengths over time to calculate different parameters of microtubule dynamics (Figure 3A). The β3lat allele displays an increase in catastrophe frequency (1.327 events/min ±0.24 compared to wild-type 0.7034 events/min ±0.14; Figure 3B; Table 2) accompanied by an increase in depolymerization rate (2.619 µm/min ±0.49 compared to wild-type; 1.865 µm/min ±0.35; Figure 3C; Table 2). These data indicate that the β3 variant residues at the lateral interface are sufficient to increase catastrophe frequency and depolymerization rate.

**Figure 3:**
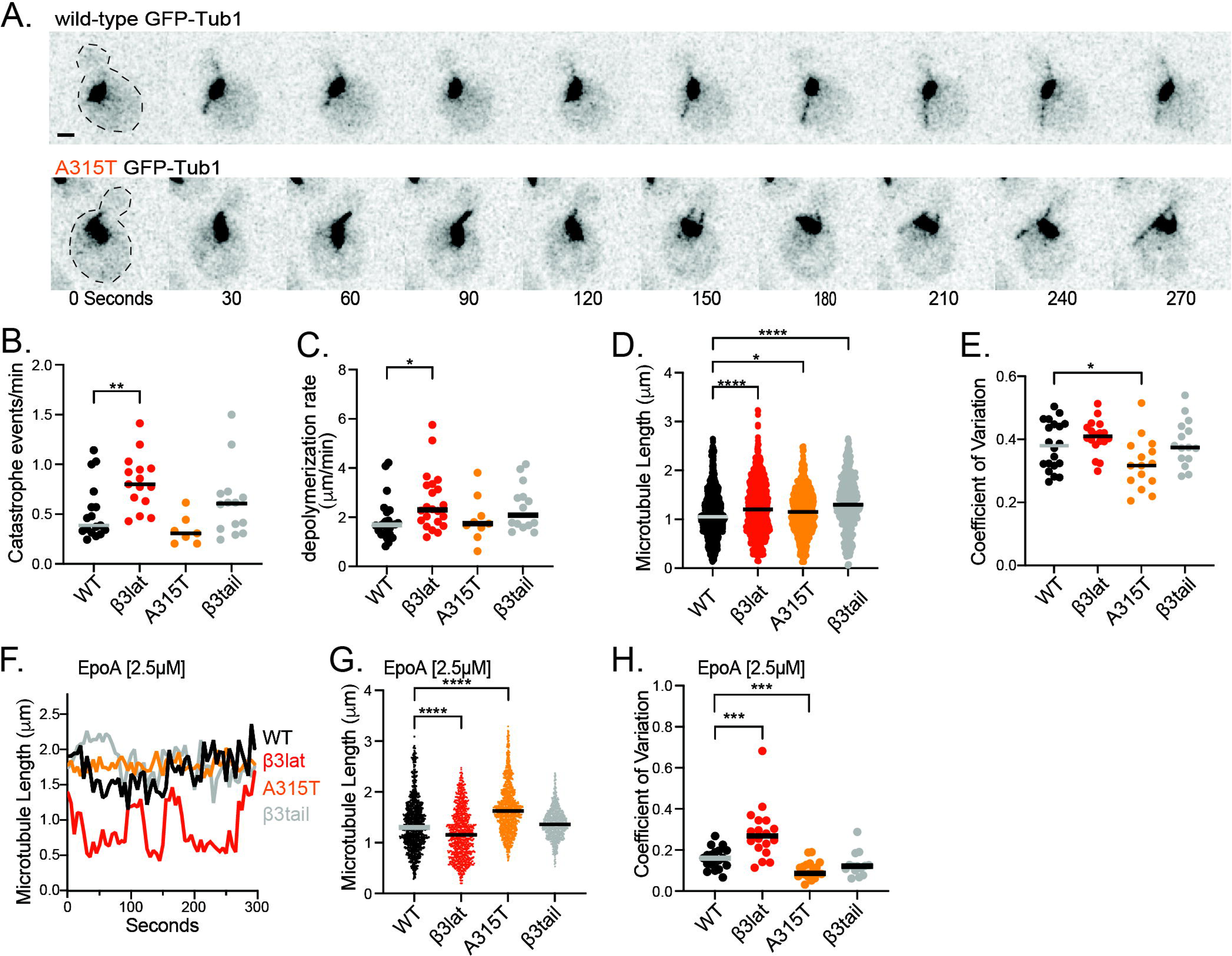
β3 variant residues are sufficient to alter microtubule dynamics. **(A)** Image montage series of wild-type and A315T cells expressing GFP-Tub1, at 30 second intervals. **(B)** Depolymerization rate (µm/min) of astral microtubules from timelapse imaging. Each dot represents a microtubule, lines indicate median values. **(C)** Catastrophe frequency (events/minute) of astral microtubules. Each dot represents a cell, lines indicate median. **(D)** Microtubule lengths (µm) of astral microtubules from timelapse imaging. Each dot represents a measured length at a single timepoint of the image series, lines indicate median. **(E)** Coefficient of variation for each microtubule analyzed, calculated by dividing the standard deviation by the mean. Each dot represents one microtubule. **(F)** Life plots of individual astral microtubules in 2.5 µM EpoA. Microtubules were measured at 5 second intervals. **(G)** Microtubule lengths (µm) of astral microtubules in 2.5 µM EpoA. Each dot represents a measured length at a single timepoint of the image series, lines indicate median. **(H)** Coefficient of variation for microtubules in cells treated with EpoA, calculated by dividing the standard deviation by the mean. Each dot represents one microtubule. For panels B-E, G and H, median values and p-values are reported in Tables 2 and S1.

**Table 2.** Microtubule dynamics in yeast cells. Values are median ± 95% confidence interval. N = number of microtubules analyzed. Values in boldface are significantly different from wild-type controls (p < 0.05).

| genotype | wild-type | wild-type | β3lat | β3lat | A315T | A315T | β3tail | β3tail |
| --- | --- | --- | --- | --- | --- | --- | --- | --- |
| 2.5 μM EpothiloneA | - | + | - | + | - | + | - | + |
| N | 20 | 19 | 18 | 18 | 15 | 21 | 15 | 14 |
| Median Length (μm) | 1.05 ± 0.03 | 1.30 ± 0.04 | **1.20 ± 0.06** | **1.15 ± 0.04** | **1.15 ± 0.05** | **1.63 ± 0.03** | **1.30 ± 0.07** | 1.36 ± 0.02 |
| Coefficient of Variation | 0.38 ± 0.06 | 0.16 ± 0.04 | 0.41 ± 0.02 | **0.27 ± 0.06** | **0.32 ± 0.05** | 0.09 ± 0.01 | 0.37 ± 0.04 | 0.12 ± 0.04 |
| Polymerization Rate (μm/min) | 1.38 ± 0.20 | --- | 1.75 ± 0.41 | --- | 1.14 ± 0.42 | --- | 1.21 ± 0.19 | --- |
| Depolymerization Rate (μm/min) | 1.69 ± 0.34 | --- | **2.30 ± 0.48** | --- | 1.74 ± 0.55 | --- | **2.10 ± 0.50** | --- |
| Catastrophe Frequency (events/min) | 0.39 ± 0.05 | --- | **0.8 ± 0.17** | --- | 0.31 ± 0.10 | --- | 0.61 ± 0.30 | --- |
| Percent Paused | 22 ± 13 | --- | 26 ± 12 | --- | 29 ± 17 | --- | 21 ± 14 | --- |

**Table 3.**
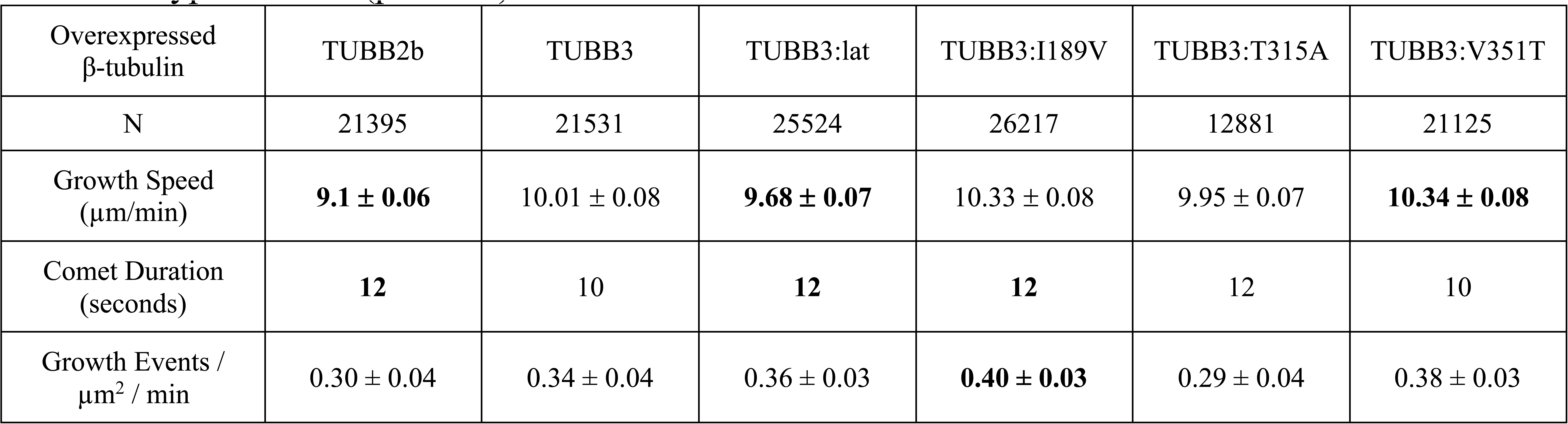
Microtubule dynamics in HeLa cells. Values are median ± 95% confidence interval. N = number of growing microtubules analyzed. Values in boldface are significantly different from wild-type controls (p < 0.05).

| Overexpressed β-tubulin | TUBB2b | TUBB3 | TUBB3:lat | TUBB3:I189V | TUBB3:T315A | TUBB3:V351T |
| --- | --- | --- | --- | --- | --- | --- |
| N | 21395 | 21531 | 25524 | 26217 | 12881 | 21125 |
| Growth Speed (μm/min) | **9.1 ± 0.06** | 10.01 ± 0.08 | **9.68 ± 0.07** | 10.33 ± 0.08 | 9.95 ± 0.07 | **10.34 ± 0.08** |
| Comet Duration (seconds) | **12** | 10 | **12** | **12** | 12 | 10 |
| Growth Events / μm<sup>2</sup> / min | 0.30 ± 0.04 | 0.34 ± 0.04 | 0.36 ± 0.03 | **0.40 ± 0.03** | 0.29 ± 0.04 | 0.38 ± 0.03 |

The A315T allele displays microtubules that curl around the cell cortex and occasionally break from the spindle pole body; reminiscent of the β3body allele (Figure 3A). A315T microtubules are slightly longer than wild-type controls and show a decreased coefficient of variation, indicative of less dynamic microtubules (Figure 3D and E; Table 2). We also noted that cells expressing A315T tended to show an increased number of astral microtubules compared to wild-type cells (data not shown).

The β3tail allele displays a modest increase in microtubule length (1.315µm ±0.05 compared to wild-type; 1.104 µm ±0.03; Figure 3D) but no change in any of the other parameters measured when compared to wild-type (Table 2). This indicates the β3 CTT allele does not strongly affect microtubule dynamics in yeast.

To test the prediction that increased microtubule dynamics in the β3lat variant persists in the presence stabilizing MTAs, we treated cells with 2.5 µM EpoA for 30 minutes and then collected time-lapse imaging of GFP-labeled microtubules. At this low concentration of EpoA, wild-type microtubules exhibit a ∼50% decrease in coefficient of variation and ∼20% increase in mean microtubule length (Table 2), compared to barely detectable length variation at higher concentrations (data not shown); therefore, 2.5 µM EpoA allows us to assay for sensitivity or resistance through changes in these parameters. When treated with 2.5 µM EpoA, microtubules in β3lat cells are significantly more dynamic than in wild-type cells or in cells expressing A315T or β3tail. This is exhibited by example lifeplots showing periods of polymerization and depolymerization for the β3lat variant, compared to relatively static lengths for the other strains (Figure 3E). Additionally, β3lat cells show a shorter mean microtubule length and increased coefficient of variation (Figure 3G and H). Overall, microtubules in cells expressing the β3lat variant are more dynamic and exhibit more length changes than wild-type controls treated with the same concentration of EpoA (Figure 3F and G). These data indicate that the variant residues at the β3 lateral interface, but not residue 315 or the CTT, are sufficient to maintain microtubule dynamics in the presence of a stabilizing drug.

### β3 requires variant residues at the lateral interface to increase catastrophe and growth rate of microtubules in HeLa cells

We next asked whether variant residues in β3 are necessary to alter microtubule dynamics in human cells expressing a mixed pool of tubulin isotypes. We sought to identify an established cell line that expresses low, endogenous levels of β3 in which we could over-express various β3 mutants and test for effects on microtubule dynamics. We compared cell lysates from a panel of immortalized cell lines and measured levels of β3 and total β-tubulin by western blotting. We find that HeLa cells express the lowest amount of β3 in relation to total β-tubulin in the cell lines we tested (Figure 4A). We therefore proceeded with HeLa cells for experiments to measure the impact of ectopically expressed β3. We created a set of plasmids to transiently transfect HeLa cells and co-express a β-tubulin under the β-actin promoter and GFP-MACF18, which tracks growing ends of microtubules. Our set of plasmids include *TUBB2b* (also highly expressed in brain tissue) as a positive control for the over-expression of β-tubulin, wild-type *TUBB3*, and *TUBB3* mutants where residues identified above as sufficient to alter microtubule dynamics in yeast are converted to mimic the β2b residue at that position. We used western blotting to confirm that each plasmid creates a 2-6 fold increase in β3 expression levels with a minimal effect on total β-tubulin expression (Figure 4B and C).

**Figure 4:**
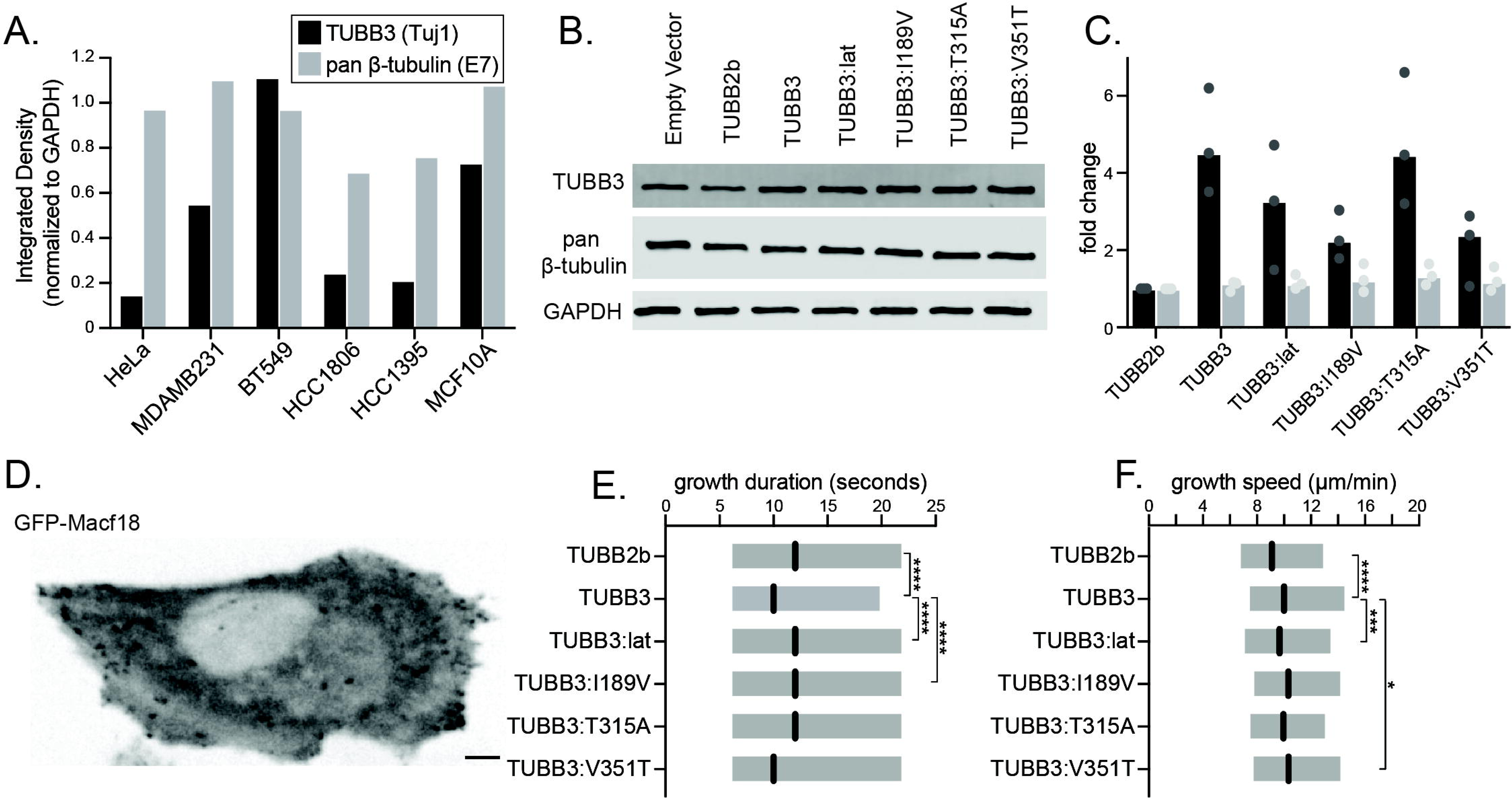
β3 over-expression alters microtubule dynamics in HeLa cells. **(A)** Quantification of β3 protein levels and total β-tubulin protein levels in indicated immortalized cell lines, based on western blotting. β3 integrated density and pan β-tubulin integrated density were normalized using GAPDH as a loading control and plotted for each indicated cell line. **(B)** Over expression of isotypes and isotype mutants in HeLa cells, measured by western blot. Control is a pCIG2 plasmid expressing GFP-MACF18 with no β-tubulin. **(C)** Quantification of β-tubulin over expression from each plasmid. β3 integrated density and pan β-tubulin integrated density were normalized using GAPDH as a loading control and plotted for each indicated overexpression plasmid. **(D)** Representative, single-timepoint image of a HeLa cell expressing GFP-MACF18. Scalebar = 5µm **(E)** Quantification of GFP-MACF18 comet duration (seconds) in cells expressing indicated β-tubulin allele. Boxes represent the 2^nd^ and 3^rd^ quartiles of the data, lines are the median values. Median values and p-values are reported in tables 3 and S2 **(F)** Comet speed (µm/minute) in cells expressing indicated β-tubulin allele. Boxes represent the 2^nd^ and 3^rd^ quartiles of the data, lines are the median values. Median values and p-values are reported in tables 3 and S2.

We predicted that increasing expression of β3 would increase catastrophe frequency in HeLa cells, and that this effect should require variant residues at the lateral interface (S33, V37, S48, S56, H57, A80, H83, L84, and I91; referred to as ‘β3:lat’) and perhaps in the core of the protein (I189, T315, and V351). To test this prediction, we imaged growing microtubule ends labeled with GFP-MACF18 in living cells that co-express the β-tubulin mutant indicated and used an automated tracking method to measure growth duration and speed. Microtubules in cells expressing β3 exhibit shorter periods of growth compared to cells expressing β2b (Figure 4E). This suggests that increasing expression of β3 increases catastrophe frequency, since GFP-MACF18 and the EB proteins to which it binds are lost from microtubule ends just prior to catastrophe (Duellberg et al., 2016). This effect of β3 requires residues at the lateral interface and the core residue V189; mutants that mimic β2b at these sites exhibit significantly longer growth durations (Figure 4E). Other β3 variant residues (T315 and V351) in the core domain do not obviously effect growth duration in this experiment.

Surprisingly, we also find that microtubule growth speed is slightly but significantly faster in cells expressing β3 compared to β2b (Figure 4F). This effect on growth speed requires the lateral interface. This suggests that in human cells, increasing expression of β3 promotes faster microtubule assembly.

### β3 variant residues at the lateral interface reduce EB1 binding and help maintain mitotic fidelity after paclitaxel treatment

Finally, we asked how increasing the proportion of β3 heterodimers in human cells alters the conformational dynamics of microtubule lattices and response to MTAs used as chemotherapeutics. To answer this question, we leveraged our GFP-MACF18 construct as a proxy for the transition zone near the end of the growing microtubule. Previous *in vitro* reconstitution studies including high-resolution cryoelectron microscopy show that EB proteins selectively bind to a transition zone just beneath the growing microtubule end, which represents an intermediate in the conversion from newly assembled GTP-bound heterodimers to the lattice of GDP-bound heterodimers (Duellberg et al., 2016, Zhang et al., 2015, Maurer et al., 2012, LaFrance et al., 2021). For growing microtubules, EB proteins exhibit a comet-like decoration of this region, with the length of the comet tail representing the rate of EB loss as heterodimers convert from the transition state to the GDP lattice state (Duellberg et al., 2016). Since MACF18 binds to EB proteins, GFP-MACF18 should exhibit a similar comet-like decoration at microtubule ends, and allow us to test whether TUBB3 accelerates the conversion of heterodimers into the GDP lattice state.

Using super resolution microscopy, we compared the localization of GFP-MACF18 at the ends of growing microtubules in live HeLa cells expressing ectopic β3 or β2b (see Materials and Methods). As expected, GFP-MACF18 exhibits a comet-like decoration of growing microtubule ends with a pronounced tail extending away from the direction of microtubule growth (Figure 5A). We measured pixel intensity values along comets and fit the tail values to a one-phase exponential decay function to calculate tail half-life and decay constant (k; Figure 5B-D). HeLa cells expressing β3 exhibit significantly shorter GFP-MACF18 comet tails with greater decay constants than cells expressing β2b (Figure 5C and D). This effect requires residues at the lateral interface of β3. These results indicate that increasing the amount of β3 in the microtubule decreases the length of the transition zone, consistent with the hypothesis that the lateral interface of β3 accelerates the conversion of heterodimers into the GDP-lattice state.

**Figure 5:**
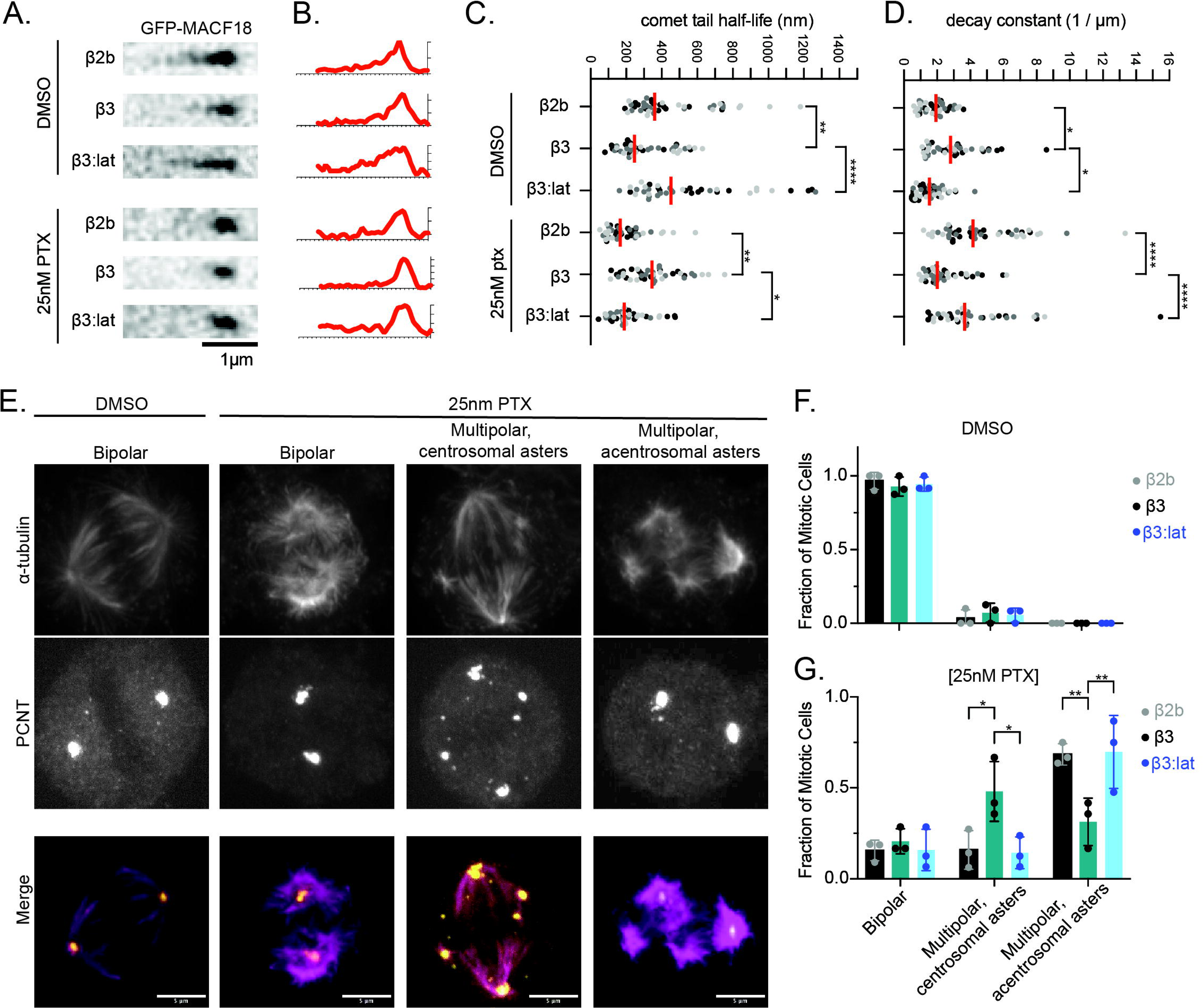
β3 over expression alters microtubule plus ends and mediates resistance to paclitaxel. **(A)** Deconvolved images of GFP-MACF18 comets collected at 280X magnification in cells expressing indicated β-tubulin allele. **(B)** Representative line scans of GFP-MACF18 comets. The x-axis represents length (100 nm intervals) and y-axis represents fluorescence intensity. **(C)** Comet tail half-life (nm) and **(D)** comet decay constant (1/µm). Dots represent individual comets, 5 collected per cell, 3 cells per experiment, collected across 3 replicate experiments. Red line represents median values for each condition. P-values are as follows, * < 0.05; ** < 0.01; ****<0.0001. **(E)** Representative images of mitotic cells for the different categories used to define spindle morphology. Cells are stained with α-tubulin and pericentrin/PCNT. **(F)** Quantification of spindle morphology in DMSO treated HeLa cells, and **(G)** HeLa cells treated with 25nM paclitaxel. Graphs represent at least 45 cells per experiment, collected across 3 replicate experiments. For each experimental replicate the fraction of mitotic cells was calculated and is represented as an individual dot on each bar. P-values are as follows, * < 0.05; ** < 0.01; ****<0.0001.

As an additional test of this hypothesis, we examined the effect of paclitaxel in cells expressing different β-tubulins. Paclitaxel binds near the lateral interface and is thought to block the conformational dynamics that normally accompany GTP hydrolysis (Alushin et al., 2014; Kellogg et al., 2017; Manka and Moores 2018; Estevez-Gallego et al., 2020). We find that cells expressing β2b exhibit a significant decrease in GFP-MACF18 comet half-life and increased decay constant when treated with 25nM paclitaxel (Figure 5A-D). These comets are also very short-lived and disappear within a few seconds (data not shown). This is consistent with paclitaxel converting heterodimers from the transition zone into an alternative state that is stable but refractory to EB binding. In contrast, GFP-MACF18 comets in cells expressing β3 are indistinguishable in paclitaxel-treatment compared to untreated controls (Figure 5A-D). Again, this effect requires the residues at the lateral interface of β3. These data are consistent with previous reports that β3 resists the effects of paclitaxel and consistent with our hypothesis that β3 promotes the conversion of heterodimers into the GDP-lattice state.

Recent studies have revealed the effectiveness of paclitaxel as a chemotherapeutic is attributable to its effects on mitosis. Cells treated with clinically relevant doses of paclitaxel form aberrant spindles that depart from the typical, bipolar configuration and exhibit declustered centrosomal components and ectopic asters of microtubules (Zasadil et al., 2014). We predicted that if β3 dampens the effects of paclitaxel on the microtubule lattice, then cells expressing increased β3 may resist the formation of aberrant spindles. To test this prediction, we stained HeLa cells expressing ectopic β-tubulins to visualize α-tubulin and pericentrin (PCNT) and assessed spindle morphology in mitotic cells. After 8 hours of treatment with 25nM paclitaxel, we find that cells commonly exhibit mitotic spindles with more than two asters of microtubules (Figure 5E). In cells expressing β2b, the ectopic, smaller asters were not associated with foci of PCNT, suggesting that these microtubules may nucleate independent of centrosomes or may detach from centrosomes after nucleation (Figure 5 E-G). In contrast, in cells expressing β3 these asters more often emanate from foci of PCNT, and the overall morphology of the spindle is more similar to the bipolar form seen in untreated cells (Figure 5 E-G). This suggests that increasing β3 expression restricts microtubule organization to centrosomes during treatment with paclitaxel. Residues at the lateral interface are required for this effect; cells expressing β3:lat show a higher frequency of aberrant spindle morphology with asters that are dissociated from PCNT foci (Figure 5G). All these data suggest that β3 promotes a lattice state that is intrinsically more dynamic and less responsive to stabilization by drugs and that the maintenance of this state may help retain microtubule organization around centrosomes during mitosis.

## Discussion

Since the discovery of tubulin isotypes almost 50 years ago, the microtubule field has sought to understand whether isotypes, particularly β-tubulins, provide redundant or distinct tubulin activities within microtubule networks. In this study, we focus on the human β3-tubulin isotype, TUBB3, to identify amino acid sequence variants that alter tubulin activity and use these variants to gain insight into how β3 distinctly affects microtubule dynamics in cells. We find that variant residues in the H1-S2 and H2-S3 loop regions of the β3 lateral interface are sufficient to increase microtubule catastrophe and promote the maintenance of dynamic microtubules in the presence of a stabilizing MTA when modeled in budding yeast β-tubulin. In human cells, we show that these residues are necessary for β3 to decrease the growth lifetime of microtubules and create a lattice state at the plus end that reduces EB binding. Furthermore, the lateral interface residues are required for β3 to resist mitotic spindle disruption by paclitaxel treatment. These findings suggest that β3 expression promotes microtubule dynamics in cells by modulating tubulin conformation during microtubule assembly and identify a central role for variant residues at the lateral interface of β3. This sheds new light on how β-tubulin isotypes alter the microtubule lattice, modulate microtubule dynamics, and change the recruitment of microtubule associated proteins (MAPs), and improves our understanding of the roles of isotypes in disease states and resistance to MTAs.

Our results show that shifting the expression of β3 alters microtubule lattice states at the growing plus end. Microtubules have long been regarded as polymers that use two states of tubulin, the GTP-cap and the GDP-lattice, to drive the behavior known as dynamic instability. However, recent work has identified additional states of tubulin between the GTP-cap and the GDP-lattice. We use the term ‘transition zone’ to represent the state (or states) that includes GDP•Pi-tubulin and exhibits the strongest affinity for EB proteins (Figure 6A; Duellberg et al., 2016, LaFrance et al., 2021). Previous reconstitution experiments with purified proteins show that the size of the transition zone increases with the speed of microtubule growth, and decreases prior to catastrophe (Zanic et al., 2009, Maurer et al., 2014, Deullberg et al., 2016). This is consistent with the transition zone representing the conversion or maturation of tubulin from the stable GTP-cap to the weaker GDP-lattice. Our results show that increasing β3 expression shrinks the transition zone in cells (Figure 6B).

**Figure 6:**
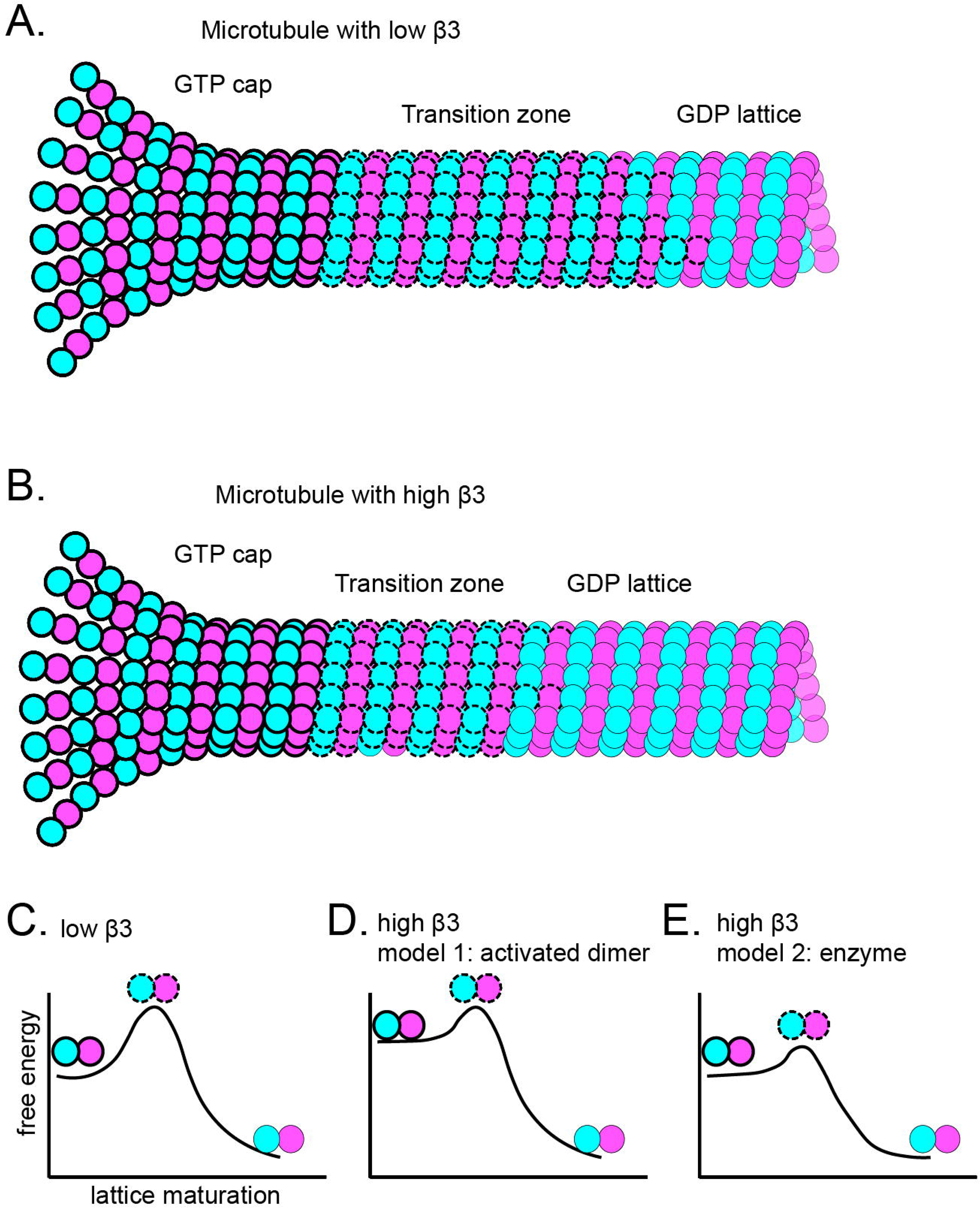
Proposed model of β3 accelerating tubulin maturation. Cartoon representation of a microtubule with (A) low β3 and (B) high β3 and the indicated lattice states. Reaction coordinate diagram models depicting microtubule lattice maturation with (C) low β3 and (D and E) high β3.

Variant residues at the β3 lateral interface could potentially impact tubulin maturation at multiple steps. During microtubule assembly, lateral interactions between heterodimers allow curved tubulin in protofilaments to straighten and completes the GTP hydrolysis site between heterodimers (Oliva et al., 2004). Within the lattice, GTP hydrolysis is accompanied by conformational changes that twist the pitch of the protofilaments and reconfigure the lateral interface (Alushin et al., 2014, Manka and Moores, 2018, Zhang et al., 2018, Zhou et al., 2023) Recent data from high-resolution structures of microtubules assembled from brain tubulin suggest that the lateral interface adopts unique conformations in the GTP-cap, transition zone, and GDP-lattice (Manka and Moores, 2018). If β3 residues in the H1-S2 and H2-S3 loops create a bias for one or more of these conformations, this could explain why we observe a shorter transition zone and more frequent catastrophes in HeLa cells expressing β3, compared to β2b (Figure 4E, 5A-D). Consistent with this possibility, a high-resolution structure of microtubules assembled from recombinant, purified β3 and stabilized with GMPCPP, a non-hydrolyzable analog of GTP, shows a displacement of the H1-S2 loop away its M loop binding partner across the lateral interface (Vemu et al., 2016). Whether β3 changes the conformation of the lateral interface for other lattice states is unknown. Importantly, our results indicate that β3 can alter lattice states even when it represents a small proportion of the β-tubulin in a cell (Figure 4B-C). This is consistent with previous studies of recombinant, purified β3 which show a non-linear relationship between the proportion of β3 and effects on catastrophe and depolymerization rate (Vemu et al., 2017, Chew and Cross, 2023). This suggests that transitions between lattice states may be highly cooperative and require the coordination of conformational changes across the lateral interface, and that β3 heterodimers alter this coordination.

If the transition zone represents a transition state of lattice maturation (Figure 6C), then β3 could lower the activation energy required for the reaction. This model is reminiscent of prior work on microtubules assembled from *C. elegans* heterodimers, which exhibit a divergent lateral interface and shorter EB comets compared to microtubules assembled from bovine brain heterodimers (Chaaban et al., 2018). β3 could similarly create ‘activated’ dimers that begin the reaction with higher free energy (Figure 6D). However, *C. elegans* dimers exhibit ∼3-fold faster assembly rates, in contrast to the slight assembly increase that we observe in cells expressing high levels of β3 (Figure 4F). This suggests that the mechanistic origins of β3 may differ from *C. elegans* tubulin. An alternative model is that β3 may lower the activation energy barrier by acting like an enzyme and reordering the lateral interface to facilitate conversion from the GTP-cap state to the GDP-lattice state (Figure 6E). We recognize that these are speculative models that will require further testing.

Could other β-tubulin isotypes exhibit β3-like properties? Our β-tubulin conservation analysis also identified TUBB1/β1 and TUBB6/β6 as divergent β-tubulins (Figure 1A-B). Both of these isotypes exhibit variant residues at the lateral interface, including several at the same positions as β3. β1 possesses 7 variant residues at the lateral interface, however only 2 of these are the same amino acid identity as β3 (Figure S1). Ectopic expression of β1 in CHO cells drives formation of curved cytoplasmic microtubules and suppresses microtubule dynamics (Yang et al., 2011). Native *TUBB1* is exclusively expressed in mature megakaryocytes and in platelet cells, where microtubules are organized into a circumferential ring called the marginal band. β6 exhibits highest sequence identity to β3 and possesses 6 variant residues at the lateral interface; 4 of these are the same amino acid identity as β3 (Figure 1A-B; S1). β6 expression increases depolymerization rate and promotes paclitaxel resistance, but the direct effect of β6 on microtubule dynamics has not been shown with *in-vitro* reconstitution studies (Bhattacharya and Cabral, 2009 and 2011). Native β6 is expressed at low levels across many tissues and cell types, but it has been associated with poor survival in several cancers (Bai et al., 2020, Li et al., 2017, Lin et al., 2019, Chung et al., 2017). Whether β6 alters conformation states in the lattice, similar to β3, will require further study.

Our findings also invite new perspectives on the mechanisms of paclitaxel action and resistance. Paclitaxel was the first microtubule stabilizing drug to be discovered and has experienced clinical success, but is met with a significant amount of resistance and therapeutic toxicity due to the broad targeting of microtubules. Paclitaxel is proposed to stabilize microtubules through binding to the M-loop of β-tubulin and stabilizing the contacts with the H1-S2 and H2-S3 loops across the lateral interface (Alushin et al., 2014, Amos et al., 1999, Li et al., 2002, Nogales et al., 1999). This locks heterodimers in an expanded state and prevents maturation. We propose that variant residues in β3 position the H1-S2 and H2-S3 loops in a way that cannot stably contact the M-loop, even in the presence of paclitaxel. Thus, β3 could resist the stabilizing activity of paclitaxel, even if the drug is bound to the M-loop of a neighboring heterodimer. Our results identifying the mechanistic origins of β3 divergence may aid in development of chemotherapeutics to more effectively target β3 and other β-tubulin isotypes.

## Methods

### β-tubulin amino-acid sequence comparison

The following accession numbers were used; TUBB1: NM030773, TUBB2a: NM001069, TUBB2b: NM178012, TUBB3: NM006086, TUBB4a: NM006087, TUBB4b: NM006088, TUBB5: NM178014, TUBB6: NM032525. *Sc*Tub2 was taken from the Saccharomyces genome database (SGD ID: SGD:S000001857). β-tubulin sequences were analyzed using Clustal Omega to create radial phylogenetic tree and NCBI Protein Blast was used to determine percent identity between pairs of sequences.

### General yeast growth and manipulation

Yeast growth, manipulation, media and transformation were performed as previously described (Amberg et. al, 2005).

### Cloning and yeast genome editing

Mutants in the budding yeast β-tubulin, *TUB2*, were generated via Quikchange site-directed mutagenesis (Agilent Technologies, Santa Clara, CA) in the *TUB2* plasmid (pJM385) and confirmed by sequencing. G126N and L170V were generated using Gibson cloning. Oligos 1883 and 1884 (G126N) and 1887 and 1888 (L170V) were used in combination with oligos 1829, 1830, 1831, and 1832 to create three fragments (around 3kb each) from pJM385 and assembled using NEBuilder HiFi DNA Assembly (New England Biolabs, Ipswich, MA). The assembled plasmid was confirmed by sequencing. A complete list of *TUB2* mutant integrating plasmids in provided in Table S4. *TUB2* coding sequence and 3’UTR containing the TRP1 marker were PCR amplified (oligos 462 and 802), gel purified and transformed into the *TUB2*/*tub2*Δ::*caURA3* strain (yJM2065). Candidate colonies were screened for loss of caURA3 and successful integration was confirmed by PCR. Diploid cells were sporulated in 1% potassium acetate and dissected to obtain haploid cells with the *tub2* mutation of interest, and confirmed by Sanger sequencing. All yeast strains are listed in Table S5.

GFP-Tub1 fusions were integrated using pSK1050 (Song and Lee, 2001) or pAFS125 (Straight et al., 1997) and expressed in addition to endogenous α-tubulin. The GFP-MACF18 construct was a gift from Casper Hoogenraad (Genentech; Cao et al., 2020). This construct was PCR amplified and placed into the pCIG2 plasmid using BsrGI and NotI sites 3’ of the IRES2 site to create pJM795. The *TUBB3* and *TUBB2b* cDNA clones were purchased from the ultimate ORF collection (IOH3755 and IOH3691 respectively; ThermoFisher Scientific; Waltham, MA). Using gateway cloning, these constructs were placed into plasmid pJM805 3’ of the β-actin promoter sequence to create dual expression plasmids with the β-tubulin of interest and GFP-MACF18. β3 mutants were generated through Quikchange of the IOH3755 cDNA clone and inserted in the pCIG2-GFP-MACF18 plasmid using Gateway cloning (ThermoFisher Scientific; Waltham, MA).

### Yeast liquid growth assay

Cells were grown to saturation in rich liquid media (2% glucose, 2% peptone, and 1% yeast extract) at 30°C and diluted 50-fold into 5 mL of fresh rich liquid media. The cultures were aliquoted into a 96-well plate, treated with 1% DMSO, Epothilone A (S1297, Selleck Chemicals, Houston, TX), or nocodazole (M1404, Sigma Aldrich, Burlington, MA) and incubated at 30°C with single orbital shaking in an Epoch2 microplate reader (BioTek; Winooski, VT). OD600 measurements were taken every 5 minutes for 24 hours. Doubling time was estimated by fitting optical density data to a non-linear logistic growth model in Prism (Graphpad, La Jolla, CA). All data gave an R^2^ value greater than 0.9 when fit to the logistic model. Each fit gave the rate constant, k, which was used to calculate doubling times using the formula t=ln(2)/k. All experiments were repeated in duplicate biological and triplicate technical replicates and wild-type controls were included in each experiment. Statistical analysis was done using a one-way ANOVA correcting for multiple comparisons with a Tukey test.

### Yeast benomyl sensitivity assay

Cells were grown to saturation in rich liquid media (2% glucose, 2% peptone, and 1% yeast extract) at 30°C and then a 10-fold dilution series was spotted onto YPD plates or YPD plates supplemented with benomyl (15 µg/mL; Sigma Aldrich #381586, St. Louis, MO). Plates were grown for two days at 30°C and then imaged.

### Microtubule dynamics in yeast

Cells were grown to log phase in rich liquid media and switched to non-fluorescent minimal media one hour before imaging. Cells were adhered to a coverslip with concanavalin A and imaged at 30°C as described in Fees et al, Jove, 2017. Briefly, Z-series spanning 6 µm with 0.45 µm steps were imaged every 5 seconds for 5 minutes. Images were collected on a Nikon Ti-E microscope equipped with a 1.45 NA 100× CFI Plan Apo objective, piezo electric stage (Physik Instrumente, Auburn, MA), spinning disk confocal scanner unit (CSU10; Yokogawa), 488-nm and 561-nm lasers (Agilent Technologies, Santa Clara, CA), and an EMCCD camera (iXon Ultra 897; Andor Technology, Belfast, UK) using NIS Elements software (Nikon). Images were maximum intensity projected using FIJI (Wayne Rasband, NIH) and lengths of astral microtubules in preanaphase cells were measured by drawing a line from the minus end, at the edge of the spindle, to the plus end. For experiments that used EpoA, the cells were gently pelleted and washed, then resuspended in non-fluorescent liquid media including 1% DMSO or EpoA at the indicated final concentration. Cells were then returned to the 30°C shaker for one hour before imaging. Statistical analysis was done using an unpaired, parametric t-test for all comparisons analyzed.

### HeLa cell culture

HeLa cells were grown at 37°C, 5% CO2, and humidity in DMEM (Gibco; Waltham, MA) supplemented with 10% FBS (Gibco; Waltham, MA) and 1% penicillin/streptomycin (Gibco; Waltham, MA).

### Western Blotting

Cells were transfected as previously described for 24 hours before collection. Cells were washed with PBS before incubating in NP-40 lysis buffer (150 mM NaCl, 1% NP-40, and 50 mM Tris-HCl) containing protease inhibitors for 5 minutes. Wells were scraped collected into a 1.5 mL microfuge tube and vortexed at 4°C for 30 minutes. Samples were pelleted at 12000 RPM, 4°C for 20 minutes and clarified lysate was transferred to a new tube containing 1:1 2.5x Lamelli buffer and boiled for 5 minutes. Samples were run on a 12% SDS-PAGE gel, transferred to a PVDF membrane and blocked for 1 hour in Intercept (PBS) Blocking Buffer (LI-CORbio; Lincoln, Nebraska) at room temperature. Membranes were probed with anti β-tubulin (E7, 1:100, DHSB, University of Iowa), anti TUBB3 (Biolegend, Tuj1;801213), anti TUBB2 (Abcam; EPR16773), and anti GAPDH (Cell Signaling Technologies; 14C10-2118) overnight at 4°C, followed by goat-anti-mouse-680 (LI-CORbio 926-68070; at 1:15000) and goat-anti-rabbit-800 (LI-CORbio 926-32211; at 1:15000) and imaged on an Odyssey Imager (LI-CORbio). Band intensities were quantified using Image Studio.

### MACF18 microtubule dynamics

Cells were plated at 5*10^4^ in a 24-well Ibidi (82426; Fitchburg, Wisconsin) glass bottom imaging plate and allowed to adhere for 24 hours. Adherent cells were transfected using the lipofectamine 3000 protocol. Briefly, lipofectamine 3000 (1 uL), p3000 reagent (1 uL), 500 ng DNA, and 50 uL opti-MEM were incubated for 20 minutes and then added to each well. Cells were imaged 24 hours after transfection. DMSO (1%) or paclitaxel (LC labs, P9600) were added at the indicated concentration one hour before imaging. MACF18 comets were imaged by acquiring a 1.5 µm z-stack every 2 seconds for 2 minutes. Images were collected on a Nikon Ti2-E inverted microscope equipped with 1.45 NA 100x CFI Plan Apo objective (Nikon Inc; Melville, NY), Nikon motorized stage, Prior NanoScan SP 600 µm nano-positioning piezo sample scanner (Prior Scientific; Rockland, MA), CSU-W1 T1 Super-Resolution spinning disk confocal, SoRa disk with 1x/2.8x/4x mag changers (3i; Denver, CO), 488 nm, 560 nm, and 647 nm laser, and a prime 95B back illuminated sCMOS camera (Teledyne Photometrics; Tuscon, AZ). Images were maximum intensity projected in ImageJ and analyzed with the Trackmate plugin (Ershov et al, 2022). Only tracks that began and ended during a images series and consisted of at least 3 frames were included in our analysis. Statistical analyses were performed using a one-way ANOVA correcting for multiple comparisons with a Tukey test.

### Super-resolution imaging and analysis of GFP-MACF18 comets

Super-resolution comets were acquired on the SoRa spinning disk confocal described above using the 100x objective with 2.8x magnification to achieve 280x magnification. A 1 µm z-stack was acquired at 0.16 µm steps every 2 seconds for 10 seconds. Images were maximum intensity projected using ImageJ and intensity profiles were created by fitting a line across the comet in the first frame and plotting intensity. Comet decay was analyzed by fitting the comet tail to a one-phase exponential decay function using Prism (Graphpad, La Jolla, CA). All curve fits gave an R^2^ value greater than 0.8. Statistical analysis was done using a one-way ANOVA correcting for multiple comparisons with a Tukey test.

### HeLa cell immunocytochemistry

HeLa cells were plated and transfected as described above and treated with 25 nM paclitaxel (LC Labs, P9600) for 8 hours prior to fixing. Cells were washed with phosphate-buffered saline (Gibco, 10010023) and then permeabilized with 0.5% Triton-X (Sigma-Aldrich, T9284). Cells were fixed with 4% PFA (Sigma-Aldrich, 158127) in PHEM buffer, which contains 60 mM PIPES (Sigma-Aldrich, P6757), 25 mM HEPES (Sigma-Aldrich, H3375), 10 mM EGTA (Sigma-Aldrich, E3889), and 2 mM MgCl2 (Acros, 223211000) for 20 min at room temperature, washed with PHEM. Cells were reduced with 10 mg/ml sodium borohydride (Fisher, S678-10) in PHEM (Acros, 177150050) for 5 min at room temperature, then washed three times with PHEM. Cells were blocked with blocking buffer, 5% BSA (Fisher, 50-253-893) in PHEM for 1 hour at room temp. Immunostaining was performed using a primary antibody directed against α-tubulin (DM1A, 1:500, Sigma-Aldrich, T6199) and pericentrin (1:2000, Abcam; Waltham, MA, ab4448). Primary antibody was diluted in blocking buffer and incubated overnight at 4°C in a humidified chamber. After primary antibody staining, cells were washed three times with PBS. The secondary antibodies goat anti-rabbit IgG Alexa Fluor 568 (Thermo, A-11011) and goat anti-mouse IgG Alexa Fluor 647 (Thermo, A32728) were diluted 1:500 in blocking buffer and incubated for 1 hr at room temperature in a dark container. Cells were sealed with glass coverslips and aqueous mounting media containing DAPI (Vector Laboratories, H-1200). Images were collected on the SoRa spinning disk confocal described above using the 40x objective with 4x magnification. Statistics represent values from a one-way ANOVA corrected for multiple comparisons by a Tukey test.

## Acknowledgements

We are grateful to members of the Moore Lab for helpful discussions. We thank Dr. Gabriela Li for help in designing the *TUB2*-integrating plasmid to create all the *TUB2* mutants, Dr. Casper Hoogenraad for the GFP-MACF18 construct, and Dr. Rytis Prekeris for the HeLa cells. This work was supported by NIH R35GM136253, CLC AWD-230491-JM, and NSF CAREER 1651841.

## Figure Legends

**Video S1:** β3full cell expressing GFP-Tub1. Video plays at 30X real time. Scale bar is 1µm.

**Video S2:** β3body cell expressing GFP-Tub1. Video plays at 30X real time. Scale bar is 1µm.

---

**Table S1.**
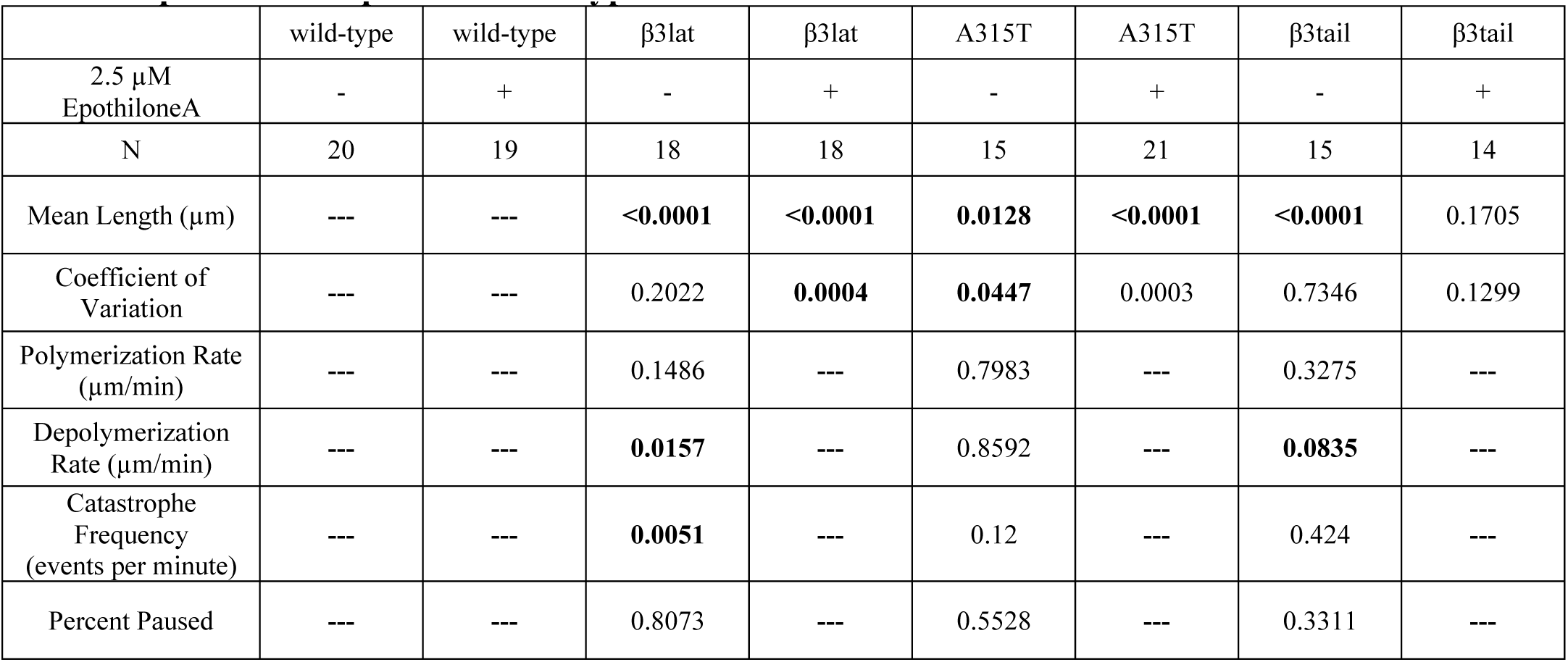
p-values compared to wild-type controls.

**Table S2.**
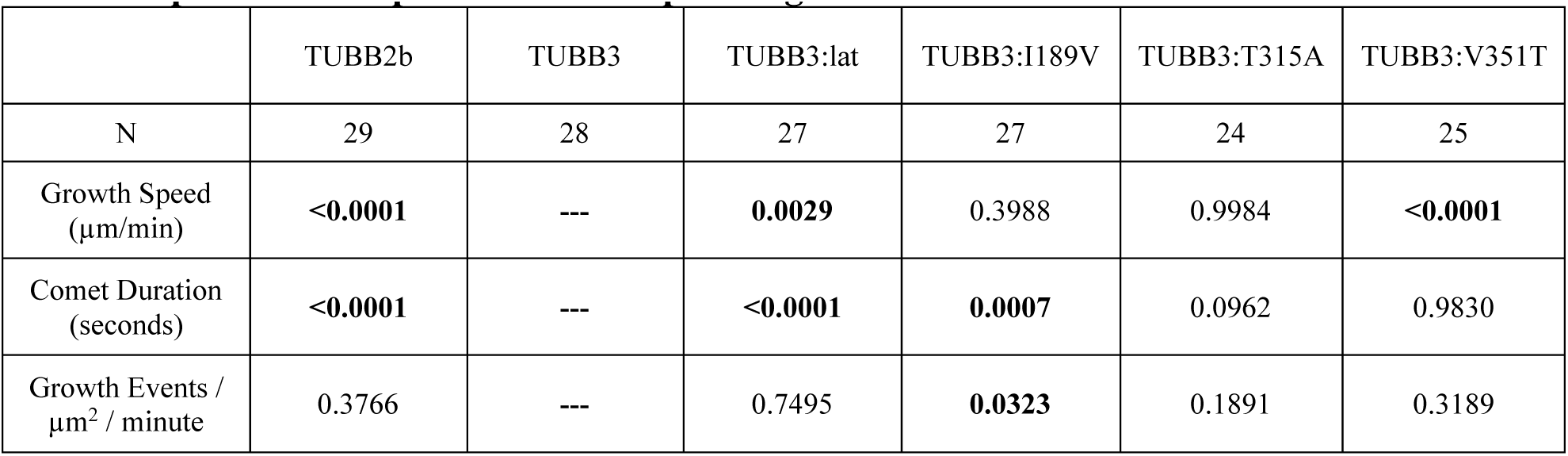
p-values compared to cells expressing TUBB3.

|  | TUBB2b | TUBB3 | TUBB3:lat | TUBB3:I189V | TUBB3:T315A | TUBB3:V351T |
| --- | --- | --- | --- | --- | --- | --- |
| N | 29 | 28 | 27 | 27 | 24 | 25 |
| Growth Speed (μm/min) | **<0.0001** | --- | **0.0029** | 0.3988 | 0.9984 | **<0.0001** |
| Comet Duration (seconds) | **<0.0001** | --- | **<0.0001** | **0.0007** | 0.0962 | 0.9830 |
| Growth Events / μm<sup>2</sup> / minute | 0.3766 | --- | 0.7495 | **0.0323** | 0.1891 | 0.3189 |

**Table S3.**
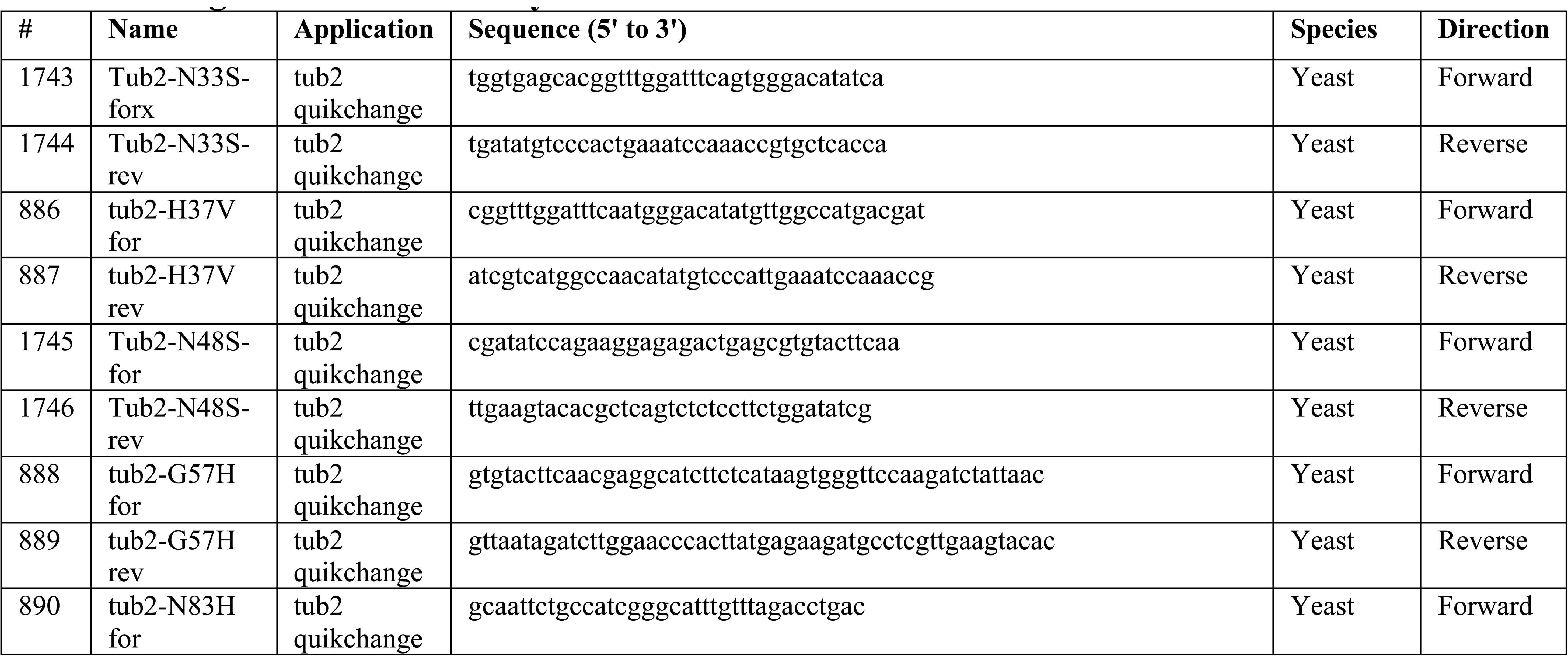
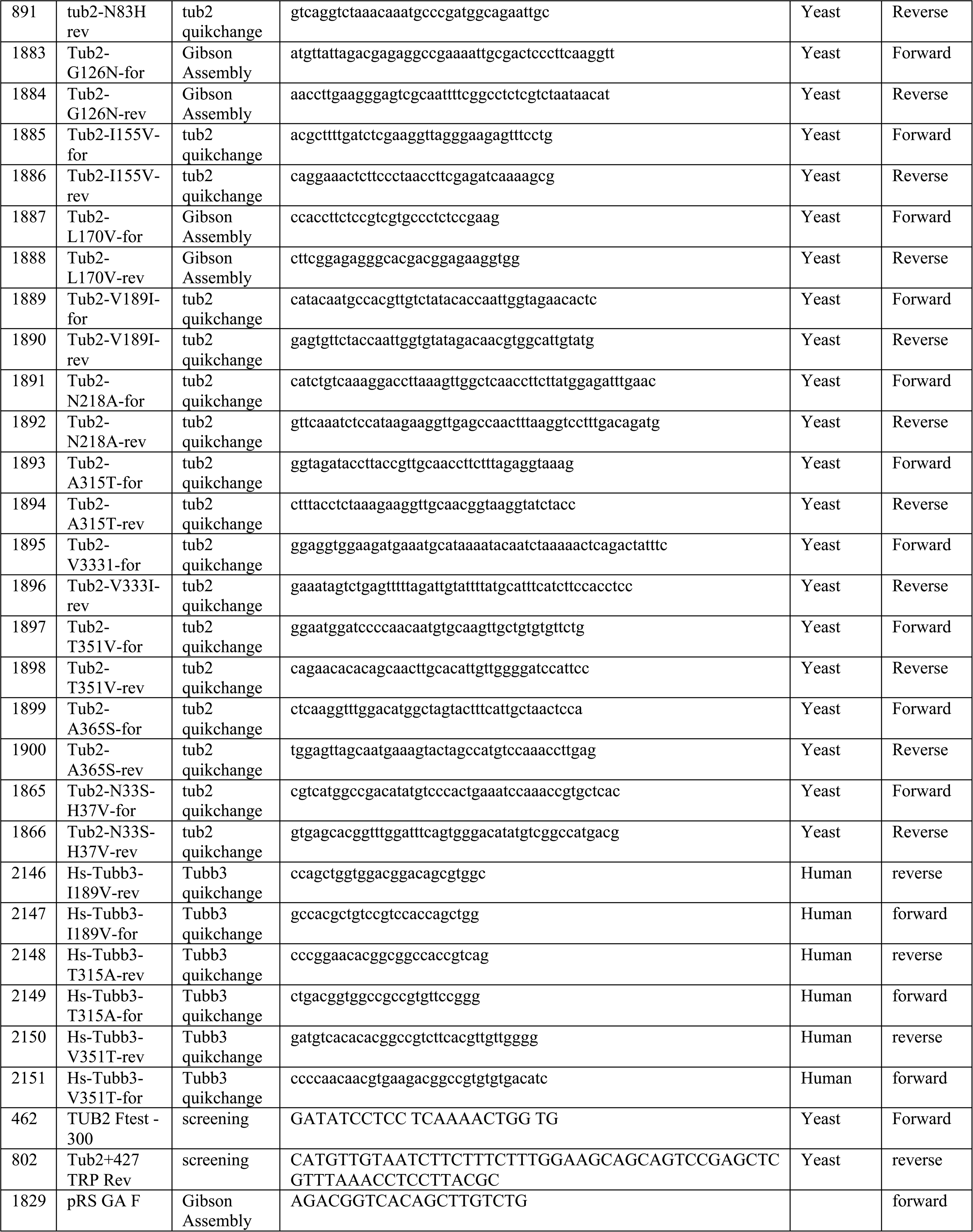
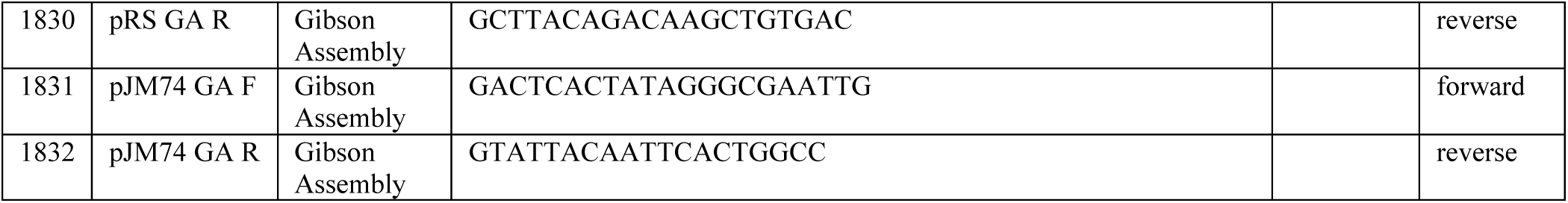
Oligos used in this study.

**Table S4.** Plasmids used in this study.

| Plasmid ID | Plasmid Name | Source |
| --- | --- | --- |
| pJM0091/pSK1050 | pUC19-LEU2::GFP-TUB1 | Song and Lee, 2001 |
| pJM0093/pAFS125 | pTUB1-GFP-TUB1-URA3 | Straight et al., 1997 |
| pJM385 | UAS-TUB2-UTR-Trp1 | This study |
| pJM576 | UAS-tub2(H37V)-UTR-Trp1 | This study |
| pJM733 | Tubb3 cDNA (IOH3755) | ThermoFisher |
| pJM734 | Tubb2b cDNA (IOH3691) | ThermoFisher |
| pJM453 | UAS-tub2(G57H)-UTR-Trp1 | This study |
| pJM454 | UAS-tub2(N83H)-UTR-Trp1 | This study |
| pJM795 | pCIG2-IRES-GFP-Macf18 | This study |
| pJM832 | pCIG2-Tubb2b-IRES-Macf18-GFP | This study |
| pJM861 | pCIG2-Tubb3-IRES-Macf18-GFP | This study |
| pJM857 | UAS-tub2(N33S)-UTR-Trp1 | This study |
| pJM858 | UAS-tub2(N48S)-UTR-Trp1 | This study |
| pJM892 | UAS-tub2(A315T)-UTR-Trp1 Iso 1 | This study |
| pJM894 | UAS-tub2(V333I)-UTR-Trp1 Iso 1 | This study |
| pJM896 | UAS-tub2(T351V)-UTR-Trp1 Iso 1 | This study |
| pJM898 | UAS-tub2(A365S)-UTR-Trp1 Iso 1 | This study |
| pJM915 | UAS-tub2(V189I)-UTR-Trp1 Iso 2 | This study |
| pJM908 | UAS-tub2(N218A)-UTR-Trp1 Iso 1 | This study |
| pJM962 | UAS-tub2(G126N)-UTR-Trp1 Iso 1 | This study |
| pJM963 | UAS-tub2(L170V)-UTR-Trp1 Iso 1 | This study |
| pJM987 | Tubb3-Tubb2lateral_pUC57-Kan | Genscript |
| pJM988 | Tubb2b-Tubb3lateral_pUC57-Kan | Genscript |
| pJM939 | UAS-tub2(N33S, H37V, N48H, G57H, N83H, G126N, I155V, V189I, N218A, A315T, V333I, T351V, A365S)-UTR-Trp1 Iso 2 | This study |
| pJM795 | pCIG2-IRES-GFP-Macf18 | This study |
| pJM832 | pCIG2-Tubb2b-IRES-GFP-Macf18 | This study |
| pJM861 | pCIG2-Tubb3-IRES- GFP-Macf18 | This study |
| pJM992 | pCIG2-Tubb3::Tubb2lateral-IRES- GFP-Macf18 | This study |
| pJM993 | pCIG2-Tubb3-I189V-IRES- GFP-Macf18 | This study |
| pJM994 | pCIG2-Tubb3-T315A-IRES- GFP-Macf18 | This study |
| pJM995 | pCIG2-Tubb3-V351T-IRES- GFP-Macf18 | This study |

**Table S5.**
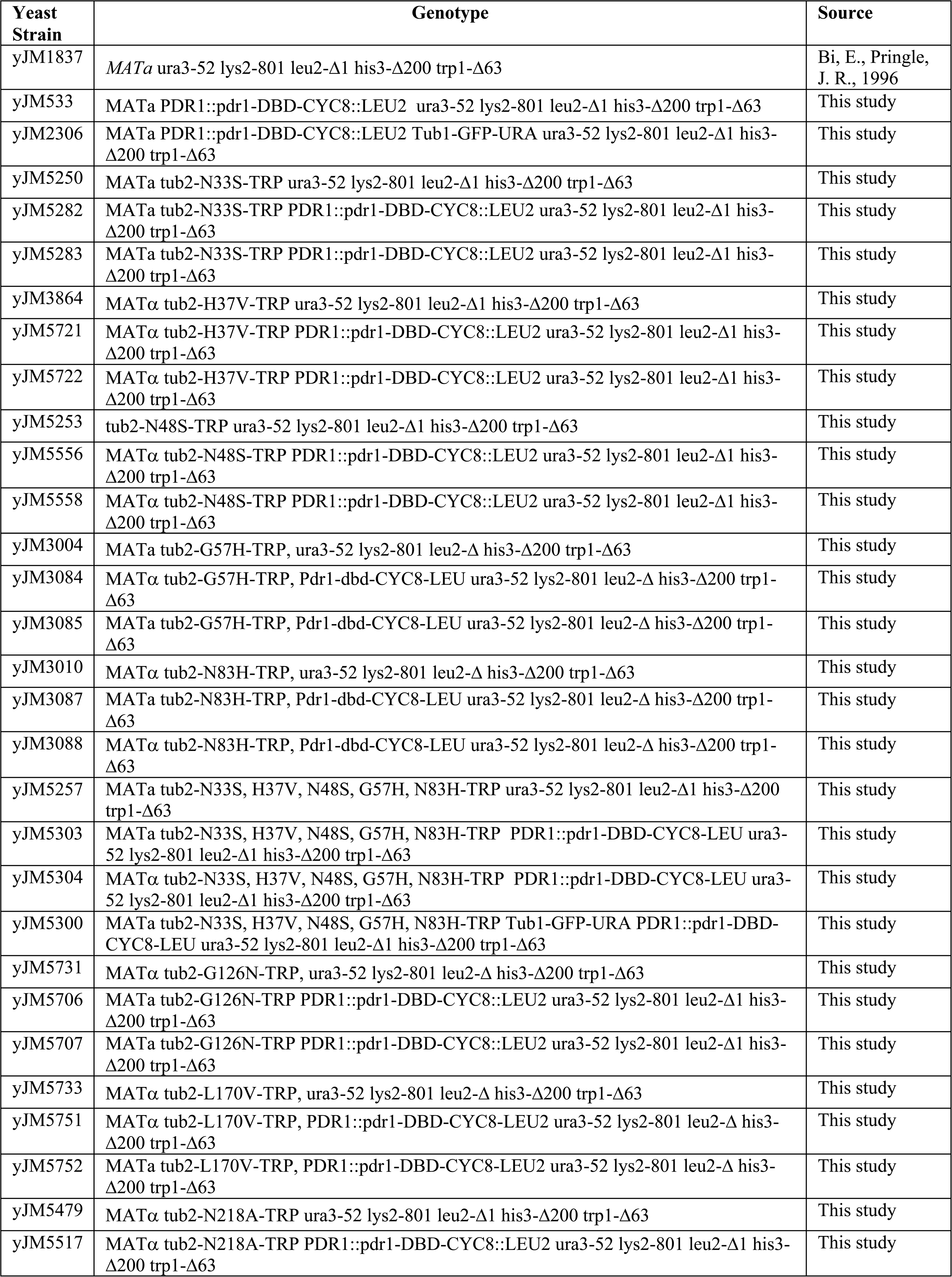
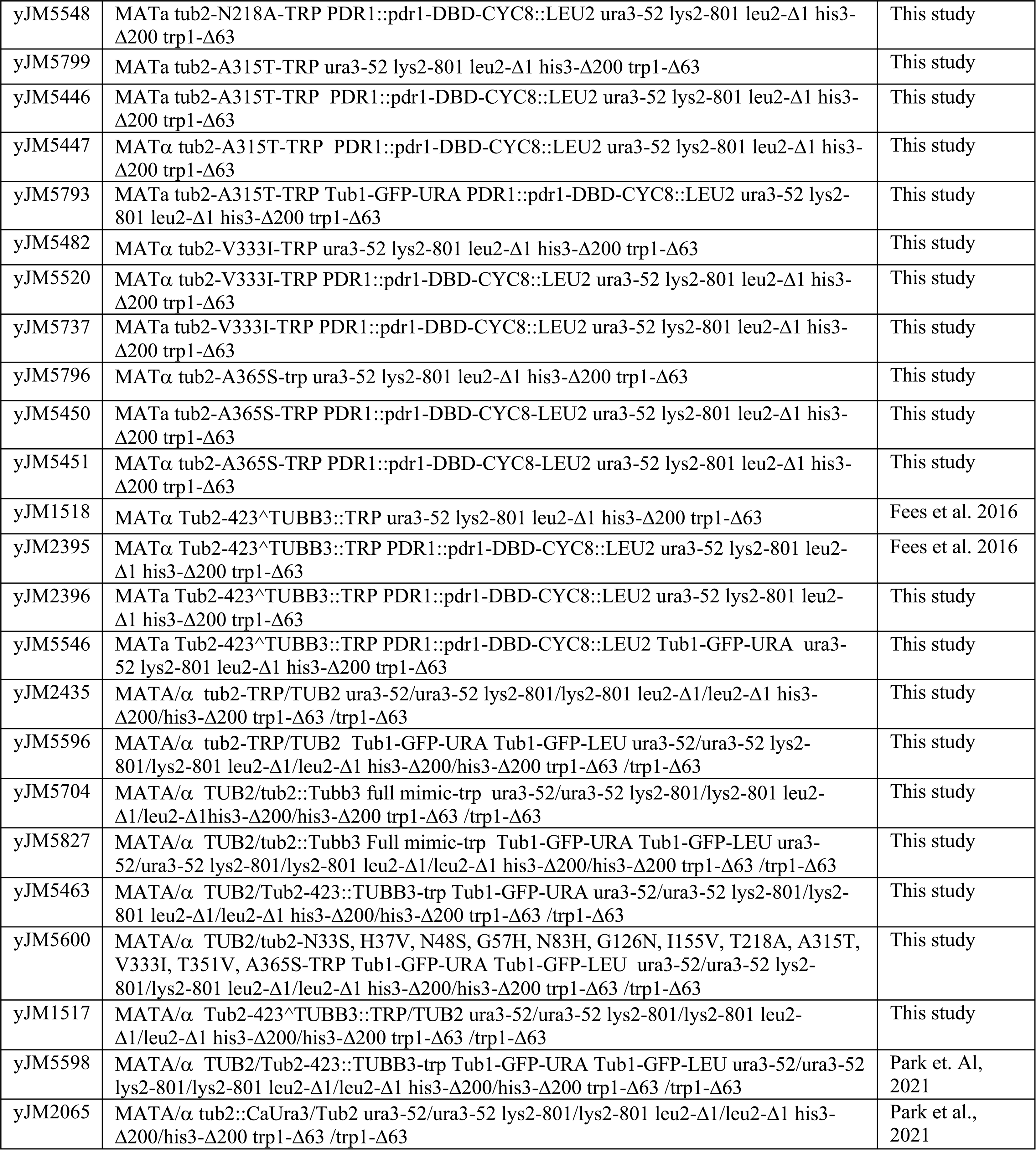
Yeast strains used in this study.

