## Supplementary figures and images for "β3 accelerates microtubule plus end maturation through a divergent lateral interface"

### Figure S1

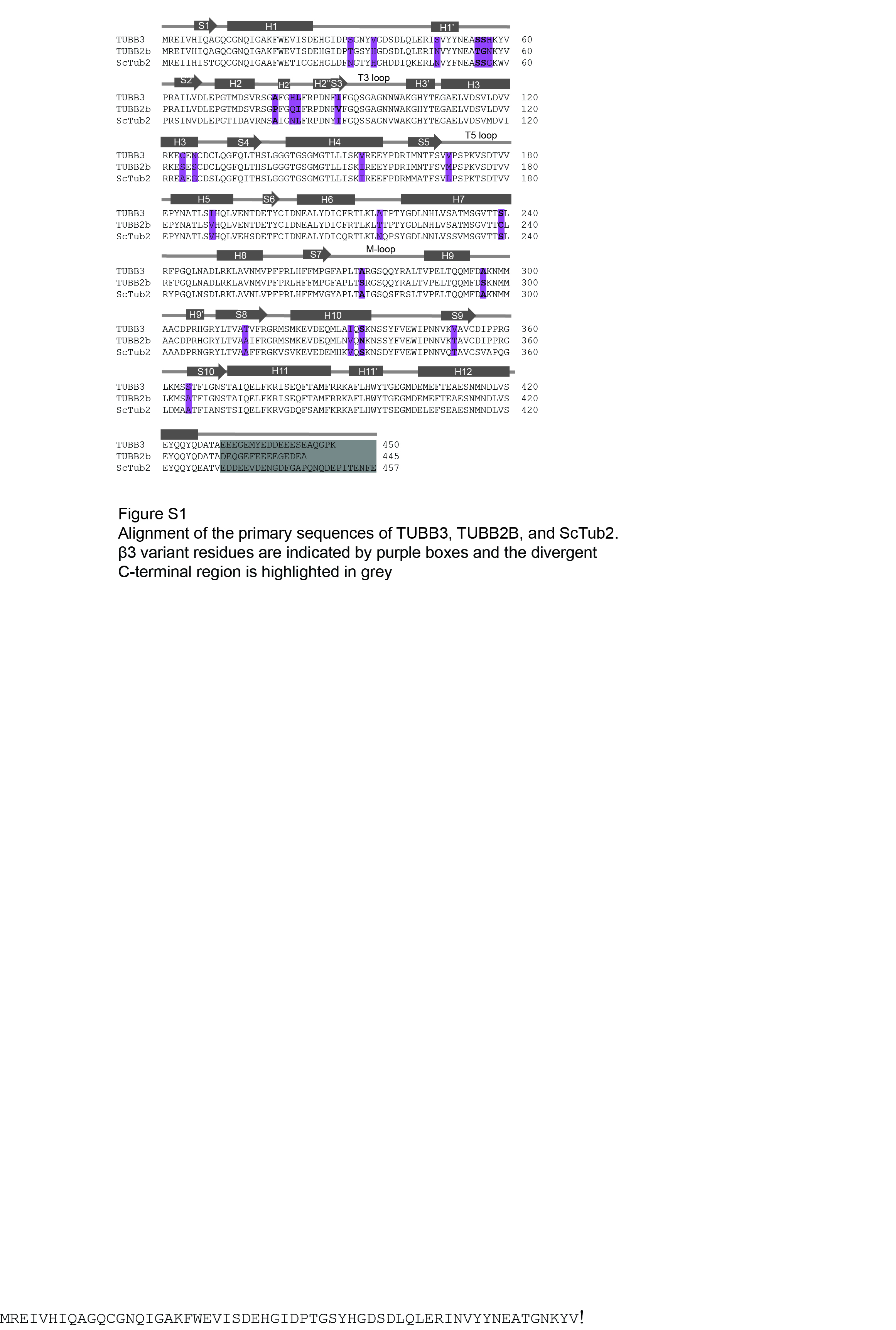

### Figure S2

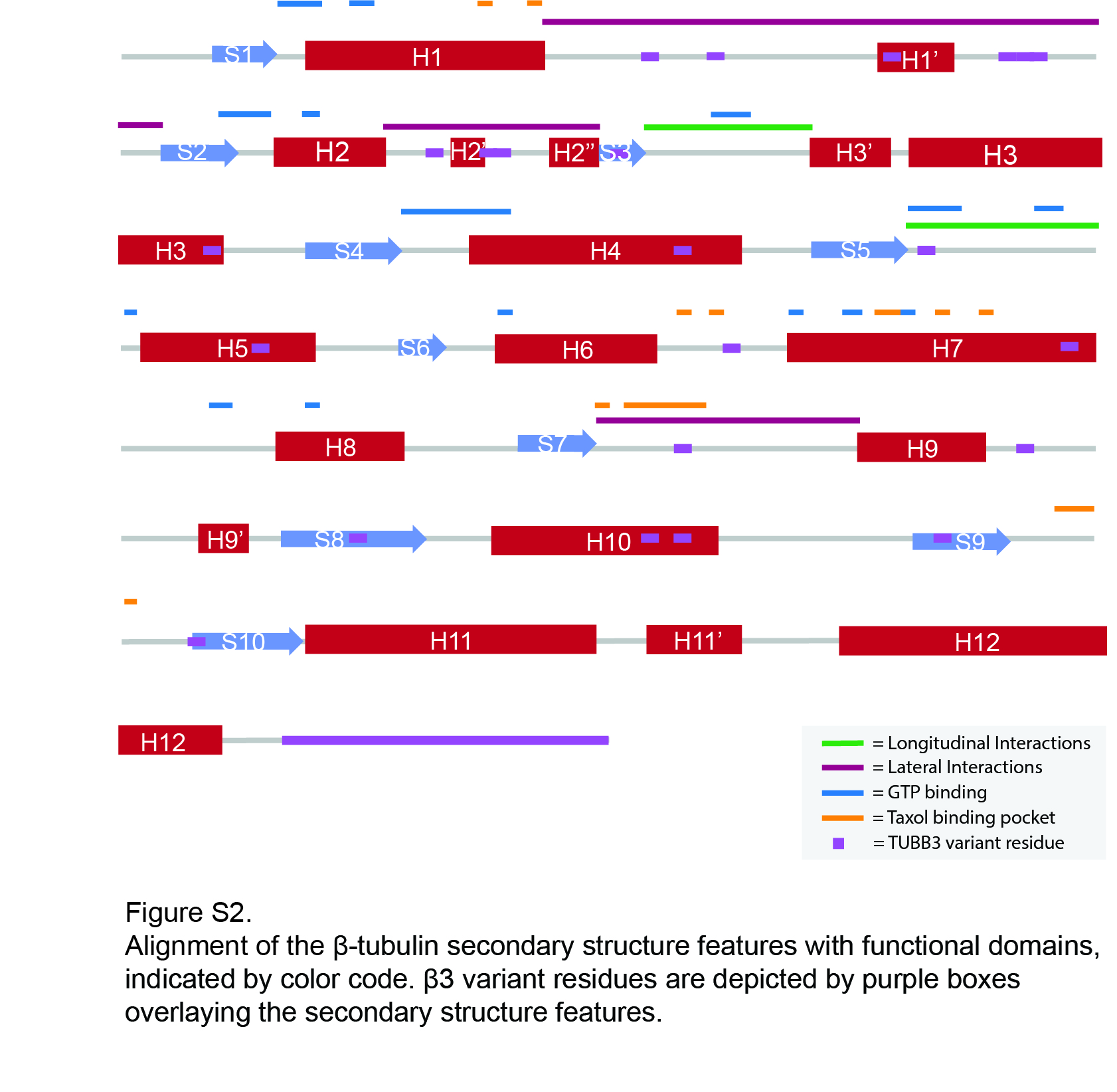

### Figure S3

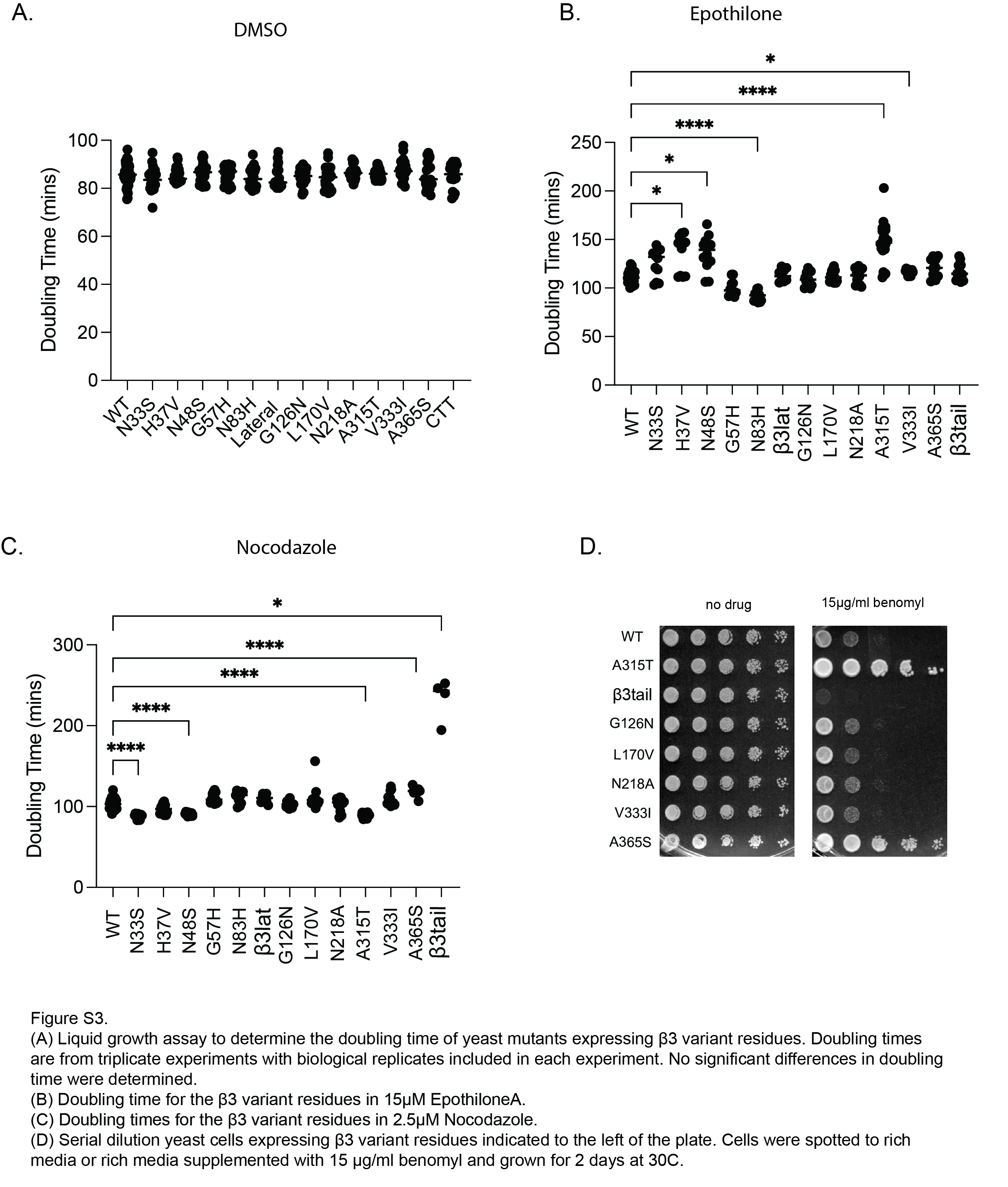
